# TorPause: Efficient Static Organ Preservation for Transplants

**DOI:** 10.64898/2026.10.08.755167

**Authors:** Joydeep Bhadury, Tetsuya Tajima, Fabian P. Suchy, Hideki Masaki, Moises Rivera, Jinyu Zhang, Toshiya Nishimura, Carsten T. Charlesworth, Carlos O. Esquivel, Hiromitsu Nakauchi

## Abstract

Transplantation is limited by how long organs survive outside the body. Current methods slow metabolism or sustain physiological support, but neither eliminates ischemia-reperfusion injury, which shortens storage and drives organ discard. Embryonic diapause and hibernation show that mammalian cells retain a latent capacity for reversible metabolic arrest. Screening the PI3K-mTOR axis, we identified a dual inhibitor that reversibly pauses embryonic stem cells at ambient oxygen and holds mouse embryos in arrest for 8.5 days. Combined with a mitochondria-targeted hydrogen sulfide donor, this yields TorPause, a two-component flush compatible with existing preservation protocols. In a rat liver transplantation model, TorPause reduced structural injury and improved recipient survival after 5 hours of static storage at room temperature, potentially expanding the usable donor pool.

## Introduction

In the U.S., more than 100,000 patients await transplantation(*1*), and roughly 17 die each day for want of a suitable organ(*2*). Many potentially usable organs are discarded because they cannot survive the interval between procurement and implantation. Static cold storage (SCS) uses hypothermia to passively depress metabolism, extending the viability window but failing to prevent early structural damage and IRI(*3–7*), while normothermic machine perfusion (NMP) provides continuous physiological support at substantial economic cost(*8*). Neither fundamentally alters the tissue’s intrinsic metabolic drive. As tissue warms, metabolic demand accelerates without matching oxygen supply, depleting ATP and priming a reperfusion burst of reactive oxygen species, calcium overload, and mitochondrial injury once flow is restored. Preventing this cascade may require actively suspending the organ’s metabolic drive rather than merely slowing it(*3, 4, 9–11*).

Nature solves this problem repeatedly. Mammalian embryonic diapause and adult hibernation are reversible metabolic pauses(*12*), and their recurrence across evolutionarily distant lineages implies conserved networks latent in vertebrates generally(*13, 14*): mammals appear to retain the machinery to pause but lack the trigger. Facultative torpor requires coordinated suppression of core body temperature, metabolic rate, and respiratory quotient(*15, 16*), of which SCS addresses only the first. The principal attempt to supply that trigger pharmacologically used inhaled H₂S, which produced an apparent suspended animation in mice(*17*). The hypometabolism, however, proved contingent on a reduced-oxygen atmosphere rather than on H₂S itself, and the approach did not extend to larger mammals, where subsequent studies found little or no hypometabolic effect(*18*).

We therefore looked instead to diapause, a conserved arrest program that had not been exploited for organ preservation. Diapause can be induced pharmacologically through mTOR inhibition in embryos, but has only been shown under 5% O₂(*19, 20*). We reasoned that this oxygen requirement reflected incomplete pathway blockade, since inhibiting mTOR alone leaves parallel PI3K signaling free to sustain residual metabolism(*21, 22*),, and that hypoxia was therefore substituting for a missing second block. Our observation that TGF-β drives mouse hematopoietic stem cells (HSCs) into a reversible hibernation-like state at 21% O₂ by suppressing PI3K-AKT signaling(*23*) gave us reason to think the pause could be uncoupled from oxygen tension altogether.

Embryonic cells are the natural substrate for this question, since diapause is their physiological state: the machinery for reversible arrest is already present and only needs the appropriate trigger. Mouse embryonic stem cells (mESCs) therefore allowed us to isolate the pharmacological switch before testing it in intact embryos and then in whole organs. A targeted screen in mESCs identified the dual PI3K/mTOR inhibitor GSK2126458 (hereafter GSK212)(*24, 25*), which we paired with the mitochondria-targeted H₂S donor AP39(*26–28*) to create TorPause. AP39 is used here not to induce hypometabolism, as systemic H₂S did, but to shield mitochondria locally during the reperfusion transition, a role that does not depend on ambient oxygen. Tor, from torpor, reflects the suspended-animation state first achieved with systemic H₂S(*17*); Pause, from diapause, denotes the GSK212-driven mTOR/PI3K pause of nutrient-sensing signaling. We show that TorPause induces reversible biostasis at ambient oxygen and extends organ viability under ambient, sub-normothermic, and hypothermic storage..

## Results

### Dual inhibition of mTOR/PI3K allows reversible pausing under normoxic conditions

To identify agents that pause mESCs at ambient oxygen (21% O₂), we screened 18 small molecules spanning the PI3K-AKT-mTOR axis: inhibitors at multiple nodes, pathway activators as controls for nonspecific perturbation, and tacrolimus, which binds FKBP12 without inhibiting mTOR, isolating FKBP12 engagement from mTOR blockade (table S1). Six mTOR-pathway inhibitors were arrayed against twelve PI3K- and related-pathway agents, applied singly or paired, each at 1 µM. Confluence (colony number × colony size, as a percentage of untreated control) was imaged longitudinally as a proxy for proliferation, and after washout to assess recovery (Fig. 1A). Cells were held throughout in N2B27 medium with LIF and 2i (PD0325901 and CHIR99021; N2B27-2iL), a defined self-renewal medium for mouse ESCs(*29, 30*), so any arrest occurred under sustained pro-proliferative signaling rather than factor withdrawal.

**Fig. 1.**
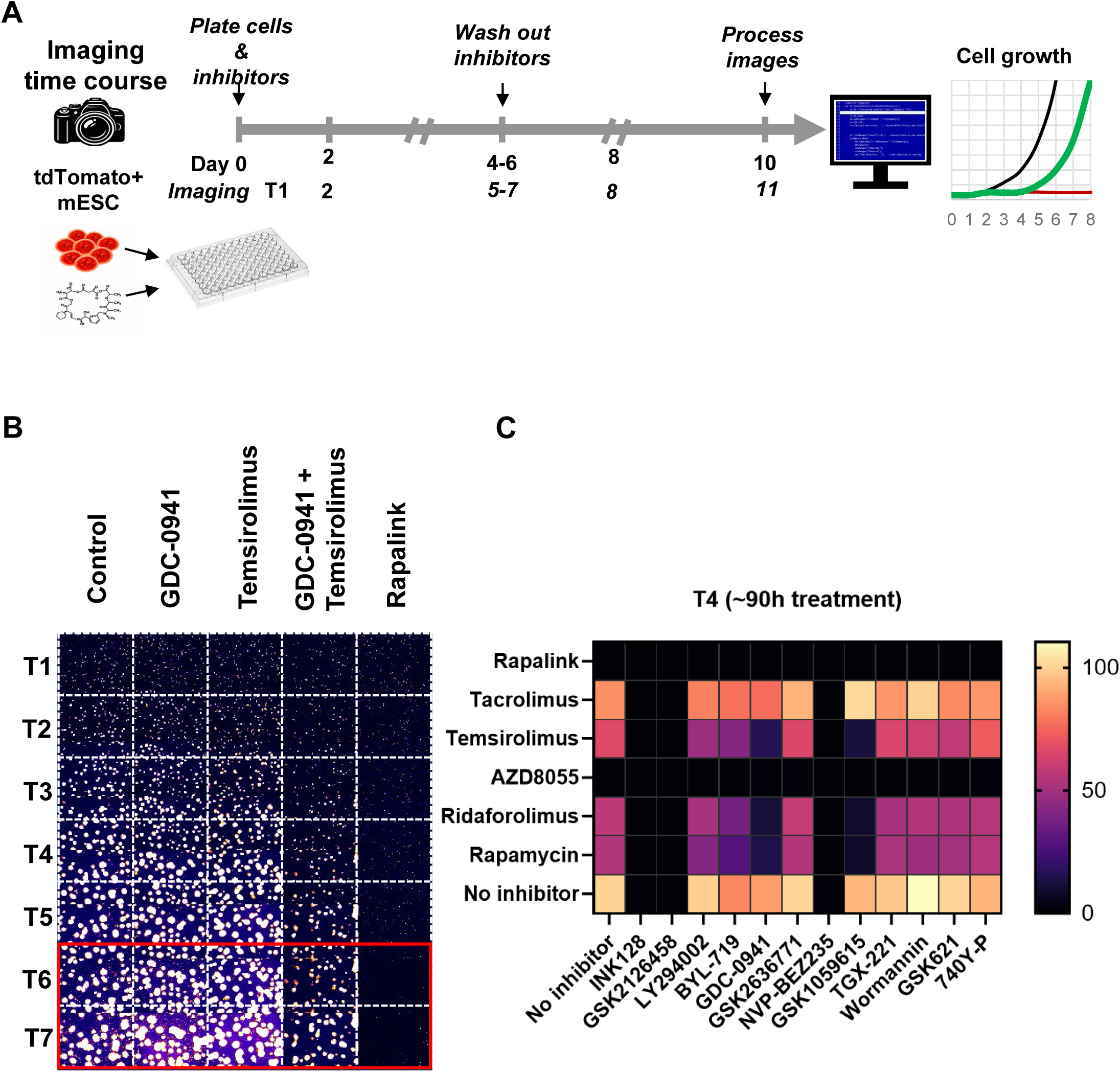
A targeted small-molecule screen identifies dual PI3K/mTOR inhibition as sufficient for reversible proliferative pausing in mESCs under normoxia. (A) Schematic of the screening pipeline. tdTomato⁺ SUN107.4 mESCs were seeded with candidate compounds and imaged longitudinally (T1 to T7) to track proliferation, followed by inhibitor washout and continued imaging to assess recovery. (B) Representative longitudinal imaging (T1 to T7) of mESC colonies treated with vehicle (Control), GDC-0941, temsirolimus, GDC-0941 + temsirolimus, or RapaLink-1. Treatment spanned T1 to T5; inhibitors were washed out after T5, and T6 and T7 are post-washout timepoints (red box). Among conditions achieving deep suppression, GDC-0941 + temsirolimus resumed proliferation after washout whereas RapaLink-1 did not recover. (C) Heatmap of cell confluence (colony number × colony size, as a percentage of untreated control) at T4 (∼90 h) across the 18 compounds screened as single agents and in pairwise combinations. Combined pan-PI3K and mTORC1 inhibition consistently outperformed single-node or isoform-selective PI3K inhibition. Compound list in table S1; full confluence data and Bliss synergy analysis in table S2, A and B.

At baseline (T1, ∼14 h) confluence was uniform across the matrix except for RapaLink-1, which suppressed growth within 12 h of seeding (Fig. 1B, fig. S1A, and table S2A). By T4 (∼90 h) clear efficacy tiers had emerged and single-node inhibition proved insufficient (Fig. 1C): untreated controls reached 100% confluence, GDC-0941 barely restricted growth (88%), and temsirolimus was only moderately effective (66%), whereas the two together suppressed confluence to 16%, far below the 58% predicted from independent action (Bliss +42).

Cooperativity depended on PI3K inhibitor class. Pairing mTORC1 inhibitors with the pan-PI3K inhibitor LY294002 gave moderate synergy (Bliss +5 to +17), and with the dual pan-PI3K/mTOR inhibitor GSK-1059615 the strongest (Bliss +42 to +49), whereas the PI3Kβ-selective TGX-221 did not exceed additivity (Bliss -0.2 to +5, within assay variability). Confluence values and synergy scores for all 72 combinations are reported in table S2, A and B.

After washout the picture inverted. GDC-0941 + temsirolimus resumed proliferation (8.7% at end of treatment, 40% by 24 h), and every combination behaved similarly, including the GSK-1059615 pairings, whereas RapaLink-1 stayed fully suppressed (≤ 0.1%) and never recovered (Fig. 1B and fig. S1, B to D). The synergy data therefore defined a requirement rather than a candidate: arrest demands blockade of both nodes. We next asked which single agents meet it in one molecule. GSK212 alone inhibits all four class I PI3K isoforms and both mTOR complexes at sub-nanomolar concentrations (table S1)(*24*), while GSK-1059615, despite its cooperativity, had negligible single-agent activity. Five compounds entered dose titration: the dual PI3K/mTOR inhibitors GSK212 and NVP-BEZ235 and the bivalent mTOR inhibitor RapaLink-1, each reaching <5% confluence as a monotherapy, plus the mTORC1/2-selective inhibitors INK128 and AZD8055, INK128 the established agent for hypoxic diapause(*19*).

#### Dose titration for reversible proliferative pausing of mESCs

To separate reversible pausing from toxicity, we titrated the five candidates across six concentrations (1 to 1000 nM) for 5 days, then washed out and tracked recovery for 6 days (N = 3; Fig. 2). Cells remained in N2B27-2iL throughout; inhibitors were removed by three washes, and proliferation was expressed as a percentage of matched 0.1% DMSO controls. Recovery was assessed both as the change in proliferation across the washout period and as the slope of a regression on recovery day, which distinguishes genuine resumption from single-timepoint fluctuation (table S3B).

**Fig. 2.**
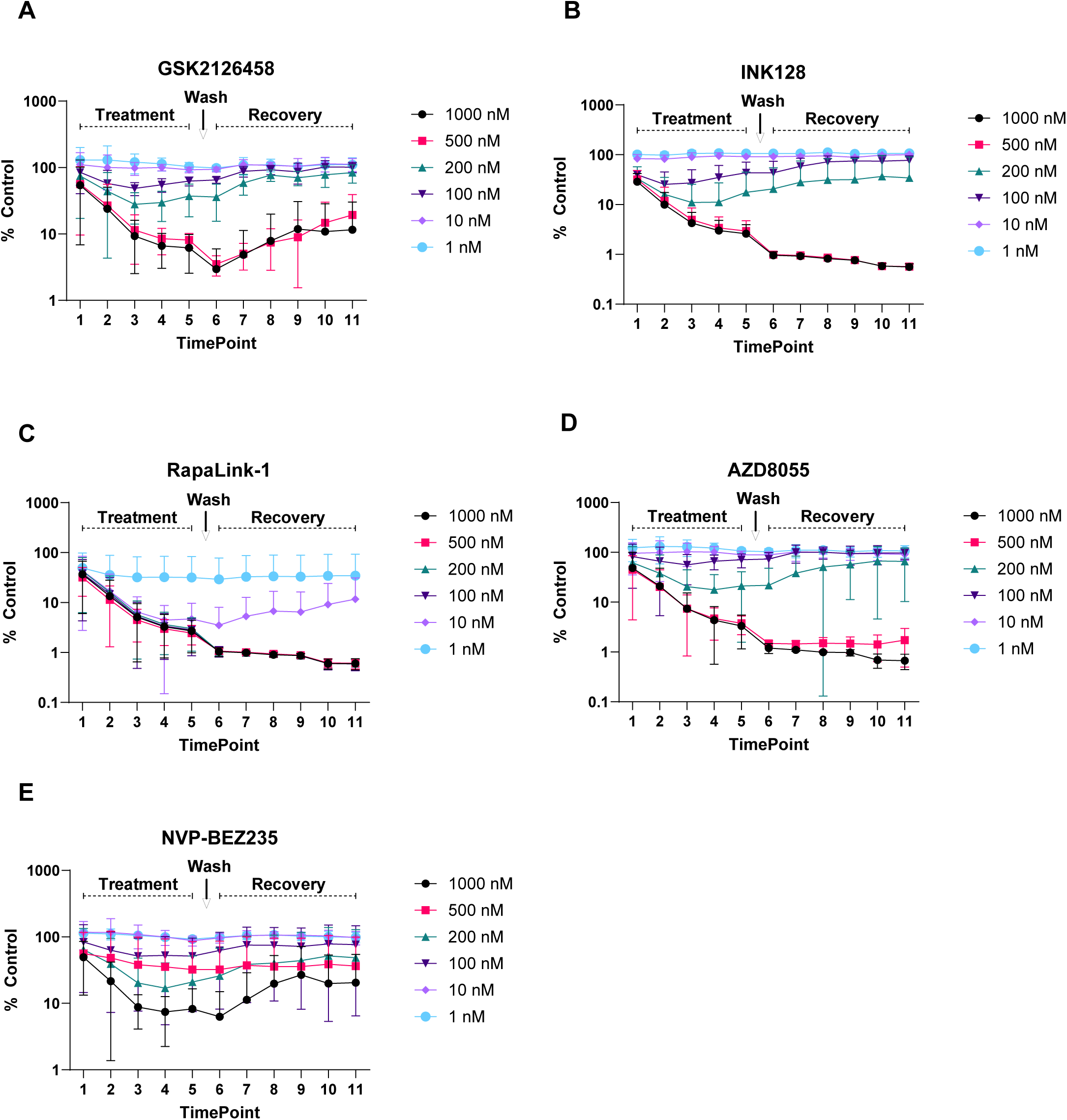
Dose titration defines the reversible pausing window and identifies GSK212 as the optimal proliferative pausing agent. Proliferation (cell confluence as a percentage of matched vehicle control) for mESCs treated with each inhibitor across a six-point concentration range (1 to 1000 nM) for 5 days (Treatment), followed by washout (arrow) and 6 days of recovery monitoring (Recovery). Data are mean ± SD of N = 3 biological replicates. (**A**) GSK212, (**B**) INK128, (**C**) RapaLink-1, (**D**) AZD8055, (**E**) NVP-BEZ235. Concentration-dependent suppression was significant for all compounds except RapaLink-1 (linear regression on log concentration; table S3A). Following washout, GSK212 uniquely recovered at two adjacent concentrations (100 and 200 nM) and was the only compound with a positive recovery slope at every concentration tested, whereas RapaLink-1 showed no recovery, NVP-BEZ235 recovered erratically, and INK128 and AZD8055 recovered only within a single concentration (recovery statistics in table S3B). Vehicle control (0.1% DMSO) did not alter proliferation kinetics.

Suppression was concentration-dependent for INK128, AZD8055, GSK212, and NVP-BEZ235 (linear regression on log concentration, all P < 0.001; table S3A) but not for RapaLink-1 (P = 0.07), which suppressed maximally at every concentration ≥ 10 nM, leaving no usable range. Washout separated the compounds sharply. GSK212 plateaued at near-maximal suppression across 500 and 1000 nM, yet cells recovered even from these concentrations, rising from 3.0% to 11.6% of control at 1000 nM and 3.5% to 19.4% at 500 nM; recovery was faster and more complete at 100 and 200 nM. GSK212 was the only compound with a positive recovery slope at every concentration tested (table S3B). For the others, maximal suppression came at the cost of recovery: INK128 and AZD8055 each recovered at a single concentration (100 and 200 nM, respectively), NVP-BEZ235 recovered erratically and without concentration dependence, all four showed negative slopes at 500 and 1000 nM, and RapaLink-1 never recovered.

#### GSK212 enables pausing of murine embryos under normoxic conditions

Although 100 nM GSK212 gave the fastest recovery in mESCs, we reasoned that intact embryos, with their greater diffusion barrier, would require more, and titrated 0.2 to 200 nM. Because spatial drift compromises long-term embryo imaging, we built an automated tracking and cropping algorithm that holds each embryo in the optical center, so the same embryo can be followed frame by frame (segmentation and tracking, fig. S2, A to C and movie S1; adaptive cropping, fig. S2D and movie S2). Using this pipeline (Fig. 3A), 2-cell embryos from naturally mated females were cultured at 37°C and 21% O₂ in medium alone or with 0.2, 2, 20, or 200 nM GSK212 or INK128, the single-node benchmark, since 200 nM is the concentration established for pausing under 5% O₂(*19*). Each embryo occupied a fixed, indexed well, so morphology was tracked in the same individuals from 1.5 to 6.5 days post-coitum (DPC) rather than sampled across embryos (Fig. 3C and movie S3). Controls progressed through normal milestones to blastocyst before degrading late in culture, whereas GSK212 held embryos in a stable pause at the blastocyst stage (Fig. 3, B and C).

**Fig. 3.**
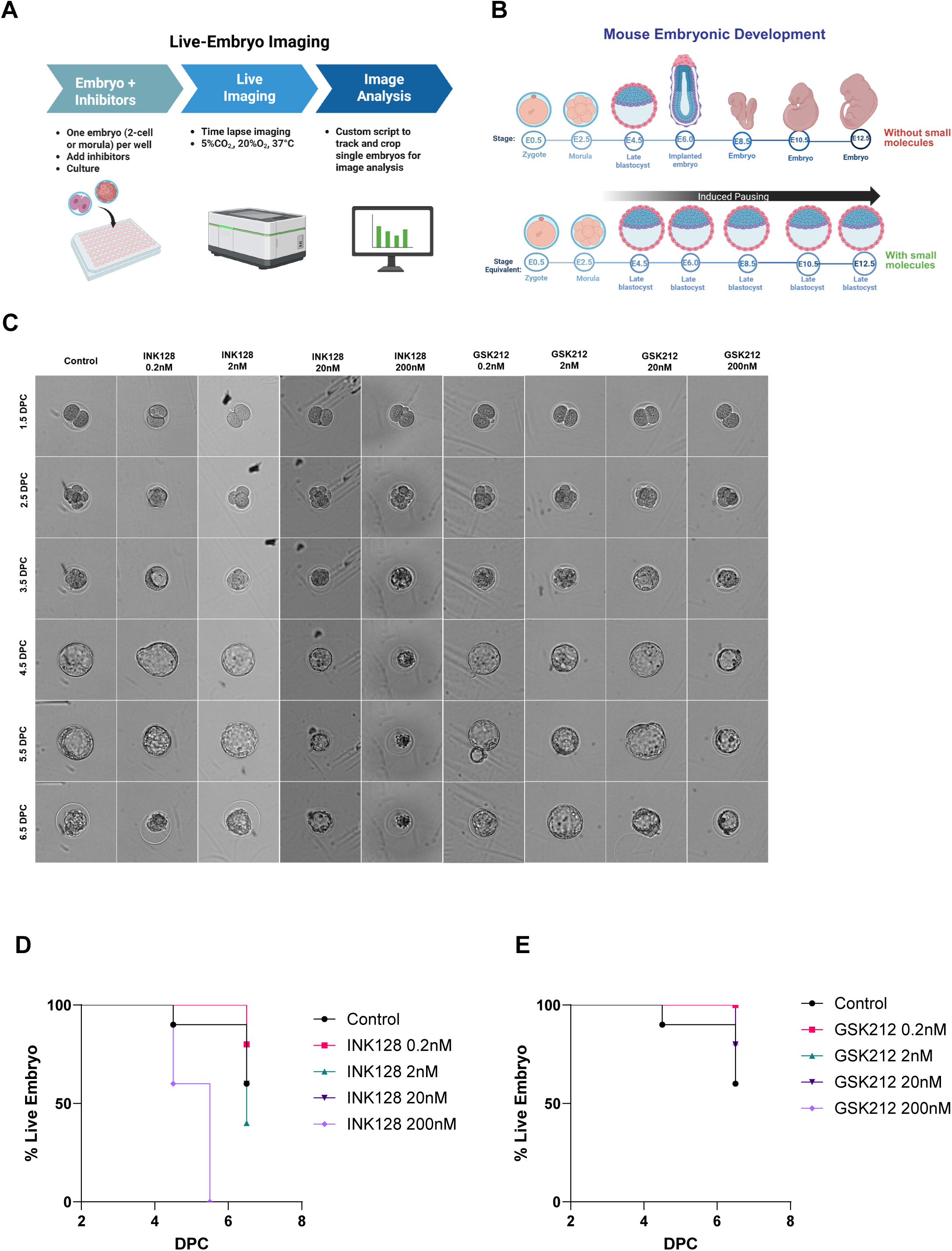
GSK212 induces a stable developmental pause and preserves viability in murine embryos. (**A**) Schematic of the live-embryo *ex vivo* pausing assay and imaging pipeline. (**B**) Diagram comparing standard *in vivo* developmental progression with the targeted GSK212-induced pause at the blastocyst stage. (**C**) Longitudinal morphological imaging of murine embryos from 1.5 to 6.5 DPC across a range of inhibitor concentrations. Each series follows the same individual embryo, imaged serially in a fixed well position; embryos were not pooled, replaced, or resampled between timepoints. Embryos were imaged at 30-min intervals and scored for viability at 24-h intervals from the complete time-lapse series (movie S3). (**D** and **E**) Kaplan-Meier survival curves for embryos cultured at 21% O₂ with vehicle (Control, n = 10) or 0.2, 2, 20, or 200 nM INK128 (**D**) or GSK212 (**E**) (n = 5 per concentration). The same control cohort is shown in both panels and is counted once in the analysis (total n = 50). Survival differed across the nine groups (log-rank χ²(8) = 52.70, *P* < 0.0001). GSK212 significantly improved survival over INK128 at 200 nM (*P* = 0.0156, Holm-corrected) but not at lower concentrations. Full pairwise comparisons in table S4A.

Tolerability diverged sharply at ambient oxygen. INK128 was toxic in a concentration-dependent manner: at 200 nM embryos fragmented and lost structural integrity, while 0.2 to 20 nM caused little acute toxicity but failed to pause development. Under 5% O₂, however, the same concentration produced a stable blastocyst-stage pause lasting more than two weeks, with 3 of 10 embryos intact and arrested at 19.5 DPC (fig. S3), consistent with pausing at reduced oxygen(*19*) and establishing that INK128 pauses embryos only when oxygen is reduced. At 21% O₂, GSK212 at 200 nM left embryos morphologically intact and viable throughout.

Kaplan-Meier analysis quantified these differences (Fig. 3, D and E). Survival differed across the nine groups (log-rank χ²(8) = 52.70, P < 0.0001). INK128 at 200 nM fell to 60% by 4.5 DPC and 0% by 5.5 DPC, whereas GSK212 at 200 nM retained 80% at the 6.5 DPC endpoint (χ² = 8.73, P = 0.0156, Holm-corrected), while the two compounds were indistinguishable at 0.2 to 20 nM (all P > 0.19, Holm-corrected; table S4A). Survival in the low-concentration arms reflected uninterrupted development rather than pausing (Fig. 3C); only 200 nM GSK212 arrested embryos at the blastocyst stage, and this concentration was carried forward.

Because most control embryos survived only about five days in vitro, we extended the time course and repeated the experiment over 8.5 days (2.5 to 11.0 DPC). Using IVF embryos for controlled developmental timing (n = 9 per condition), we compared 200 nM GSK212 against 200 nM INK128 and untreated controls under the same pipeline. Controls developed normally before succumbing to culture limitations, degrading completely by 7.5 DPC. As in the titration, INK128 was rapidly toxic, fragmenting most embryos by 4.5 DPC with none surviving past 6.5 DPC. GSK212 held a stable pause, with architecture intact through 11.0 DPC (Fig. 4A and movie S4).

**Fig. 4.**
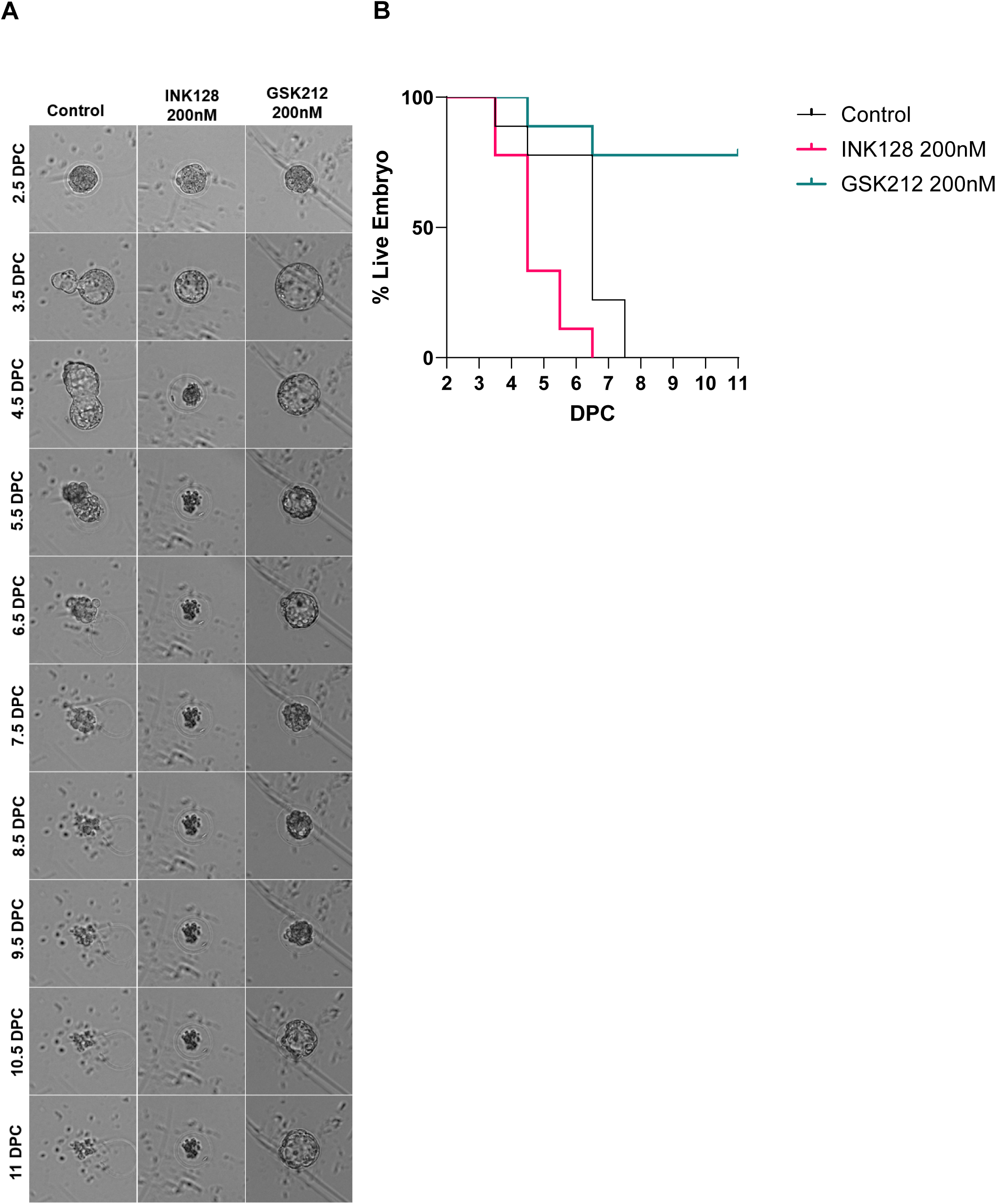
GSK212 sustains prolonged *ex vivo* metabolic pausing in IVF embryos. (**A**) Longitudinal morphological tracking of IVF murine embryos from 2.5 to 11.0 DPC. Each series follows the same individual embryo, imaged serially in a fixed well position; embryos were not pooled, replaced, or resampled between timepoints. Untreated controls degraded by 7.5 DPC. INK128 (200 nM), the concentration established for pausing under hypoxia, caused rapid toxicity and fragmentation by 4.5 DPC at 21% O₂, whereas GSK212 (200 nM) preserved intact blastocyst architecture through 11.0 DPC. Embryos were imaged at 3-h intervals and scored for viability at 24-h intervals from the complete time-lapse series (movie S4). (**B**) Kaplan-Meier survival curves for embryos cultured at 21% O₂ for up to 11.0 DPC with vehicle (Control), INK128 (200 nM), or GSK212 (200 nM) (n = 9 per group). Survival differed across the three groups (log-rank χ²(2) = 20.69, *P* = 3.2 × 10⁻). GSK212 significantly extended survival relative to both control (*P* = 0.0043) and INK128 (*P* = 0.0008), and INK128 reduced survival relative to control (*P* = 0.0086; all Holm-corrected). 78% (7 of 9) of GSK212-treated embryos remained viable at the endpoint. Full pairwise comparisons in table S5A.

Survival differed across the three groups (log-rank χ² (2) = 20.69, P = 3.2×10^-5^; Fig. 4B and table S5A). Both INK128 and untreated controls fared significantly worse than GSK212 (χ² = 13.38, P = 0.0008 and χ² = 9.40, P = 0.0043, respectively, Holm-corrected), and 78% of GSK212-treated embryos (7 of 9) remained viable at 11.0 DPC. Notably, INK128 at its hypoxic pausing concentration was not merely ineffective at ambient oxygen but actively deleterious, killing embryos faster than no treatment, leaving GSK212 as the only agent capable of sustaining embryonic pausing at 21% O₂.

#### Pharmacological pausing preserves cellular viability at sub-normothermic temperature

Natural diapause operates near body temperature (37°C). To ask whether pharmacological pausing works below normothermia, we stored an independent mESC line at room temperature (23 ± 2°C, RT), storing 1×10 cells per cryovial for three days in maintenance medium with or without 200 nM GSK212. At RT, GSK212 preserved both viability and absolute live-cell recovery across all three replicates (fig. S4A). The effect was temperature-dependent: at 4°C it conferred no benefit, with recovery matching or falling below control (fig. S4B), consistent with deep cold already suppressing the metabolic activity on which the pause acts. That GSK212 protected cells at RT prompted us to ask whether it could preserve intact organs.

### GSK212 and AP39 combination treatment mitigates ischemia-reperfusion injury during ambient liver storage

Historical attempts to induce suspended animation with H₂S depended strictly on reduced O_2_. We reasoned that pairing H₂S donor AP39 with GSK212’s oxygen-independent pause would remove that constraint. RT static storage imposes a stringent, clinically relevant challenge, since warm ischemia drives injury across solid-organ transplantation(*31, 32*): tolerable WIT is ∼30 minutes for human hepatic(*33*), renal(*34*), and cardiac(*35*) grafts and ∼60 minutes for lungs, which tolerate more owing to lower metabolic demand and residual alveolar oxygen(*36*). Given the two- to three-fold higher basal metabolic rate of rodents, 5 hours of static storage at ambient temperature, followed by up to 16 min of implantation before reperfusion, far exceeds any clinically tolerated warm-ischemic exposure. Organs were flushed and stored in RT University of Wisconsin (UW) solution containing vehicle (0.1% DMSO), 3 µM AP39, 2 µM GSK212 or both. Concentrations selected empirically from the dose-ranging below (figs. S9 and S10); they exceed those in prior static-immersion AP39 studies(*37–39*), consistent with the brief single-pass exposure of a vascular flush and the greater metabolic demand at room temperature. We first asked whether the combination engages the intended biology, profiling whole organs transcriptionally before testing function in a transplantation model.

We profiled whole organs by bulk RNA-sequencing. Donor mice were perfused with UW solution containing vehicle, AP39, GSK212, or the combination, and heart, lung, liver, and kidney were harvested after 5 hours of RT storage, before reperfusion, alongside flushed, non-stored (Sham) organs (fig. S5A). These four are the most frequently transplanted solid organs(*2*), and tissue was taken at the end of storage rather than after reperfusion so the measured state reflected the stored graft itself rather than the injury response to reperfusion. Samples separated primarily by organ of origin (fig. S5B). Gene-set enrichment analysis against the within-organ Sham baseline (fig. S5C) showed that storage alone reshaped the transcriptome: untreated controls already suppressed interferon-α responses in all four organs (NES -1.7 to -2.9, FDR ≤0.01) and interferon-γ responses in liver, kidney, and heart (FDR ≤0.04). This indicates that ischemic stress itself imposes a broad transcriptional shutdown, against which any treatment-specific effect must be judged. GSK212 and the combination deepened this in the kidney, liver, and lung, adding little in the heart, where control suppression was already near-maximal. The allograft-rejection signature was the clearest treatment-specific effect, unchanged or elevated in controls (lung NES +1.6, FDR = 0.008) and significantly suppressed only by GSK212 and the combination (kidney −2.3 and −2.7; lung −1.7 and −1.6; all FDR <0.02), with a concordant but non-significant shift in liver. Because it moves in the opposite direction in controls, this suppression cannot be attributed to storage and instead tracks GSK212 exposure. AP39 alone changed little. At gene level this tracked suppression of antigen-processing (*Psmb8*, *Psmb9*, *Tap1*), interferon-regulatory (*Nlrc5*, *Irf7*, *Gbp3*), and rejection-associated transcripts (fig. S6A). Metabolic programs, including oxidative phosphorylation, mTORC1 signaling, and glycolysis, were suppressed predominantly in the liver (figs. S6B and S7B), while proliferative programs changed modestly and variably (fig. S7A; full statistics and leading-edge genes in table S7). This dampening of immune and metabolic programs is consistent with a torpor-like quiescent state resembling the immune quiescence of natural hibernation(*40*), and motivated functional testing in the liver.

#### Rat liver transplantation

Because the transcriptional signatures were obtained at the end of storage in a non-transplant preparation, they describe the state of the stored graft rather than its functional outcome. We therefore tested the combination in a rat orthotopic liver transplantation model (Fig. 5A), evaluating all initial cohorts at 2 hours post-transplantation to capture early IRI events. Macroscopic assessment revealed divergent graft quality (fig. S8): control livers showed severe congestion and dark discoloration that persisted rather than resolving across the 2-hour window, indicating failure to establish uniform perfusion rather than transient post-reperfusion hyperemia, whereas combination-treated livers recovered color and closely resembled healthy donor tissue.

**Fig. 5.**
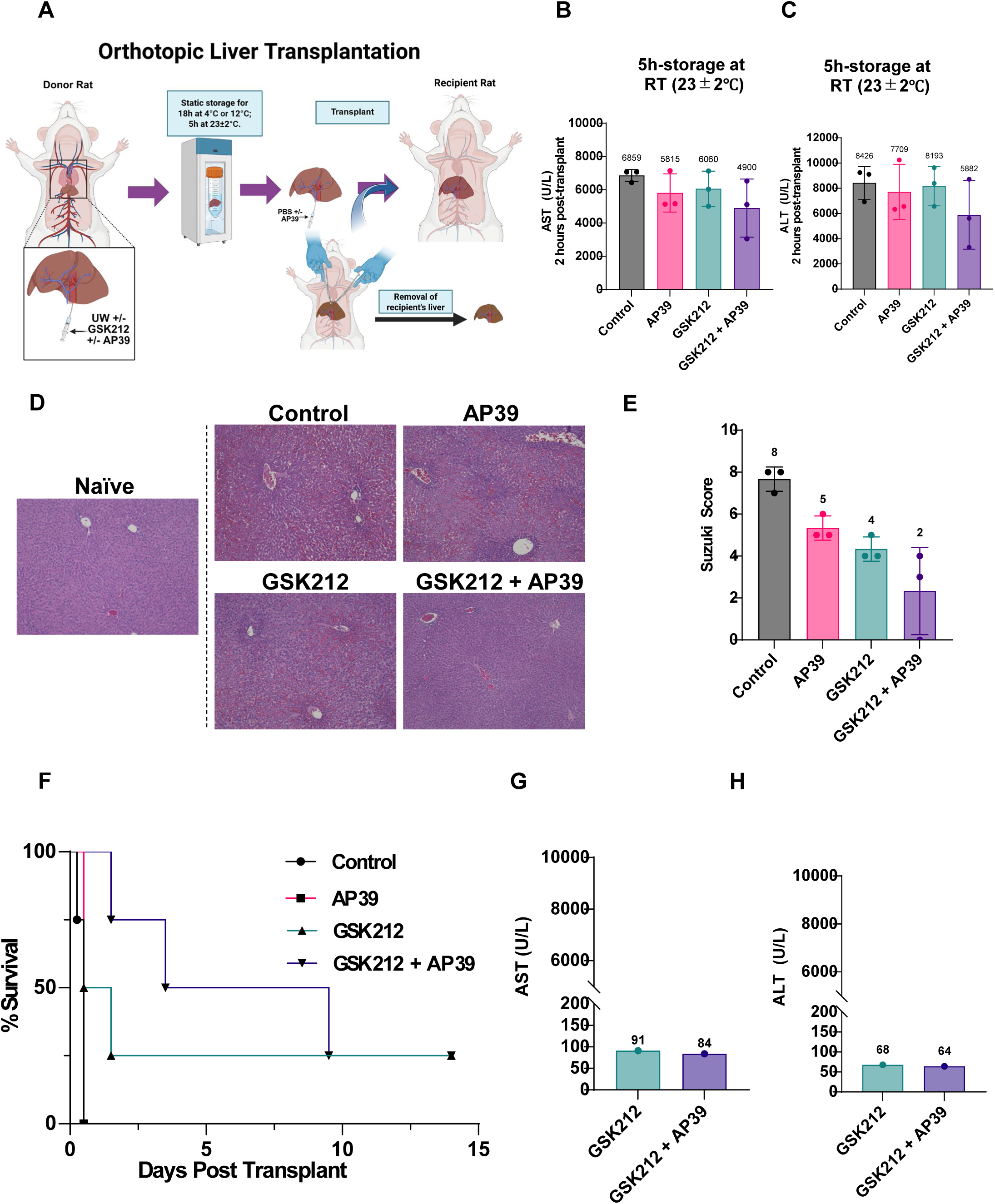
Combined GSK212 and AP39 treatment mitigates early ischemia-reperfusion injury and supports long-term survival after ambient liver storage. (**A**) Schematic of the orthotopic transplantation pipeline. Livers were flushed with University of Wisconsin (UW) solution ± GSK212 ± AP39 and held in static storage at 23 ± 2°C for 5 h, measured from interruption of donor hepatic blood flow to removal from storage for implantation; reperfusion was achieved within 16 min of clamping. (**B** and **C**) Circulating AST (**B**) and ALT (**C**) at 2 hours post-transplantation (n = 3 per group). The combination gave the lowest mean transaminases, though differences across groups were not significant (table S8). (**D**) Representative H&E-stained sections. Naïve denotes non transplanted liver after UW flush only, shown as baseline; remaining panels are grafts at 2 hours post-transplantation. (**E**) Modified Suzuki score (n = 3 per group). Combination treatment significantly reduced injury relative to control (mean 2 versus 8), as did GSK212 monotherapy; the combination did not significantly exceed either monotherapy at this sample size (table S8). (**F**) Kaplan-Meier survival over 14 days in a parallel cohort (n = 4 per group). Control and AP39 cohorts showed total lethality at the first postoperative check (0.5 days). GSK212 and combination cohorts each reached 25% survival at day 14. A Cox model confirmed a significant main effect of GSK212 on mortality hazard, with no effect of AP39 and no interaction (table S10, A and B). (**G** and **H**) AST (**G**) and ALT (**H**) in the single surviving recipient of each cohort at day 14 (n = 1 per group), both within the physiological reference range.

At 2 hours, untreated controls showed extensive injury (mean AST 6859 U/L, ALT 8426 U/L). Both monotherapies numerically reduced transaminase release, and the combination gave the lowest means (AST 4900 U/L, ALT 5882 U/L; Fig. 5, B and C), though group differences were not significant (one-way ANOVA, AST P = 0.32, ALT P = 0.45; n = 3 per group; table S8). Histology discriminated between groups more sharply than biochemistry (Fig. 5, D and E). Against naïve liver (Fig. 5D), control grafts showed severe sinusoidal congestion, vacuolization, and necrosis (mean modified Suzuki score 8); monotherapies reduced this (5 for AP39, 4 for GSK212), and the combination best preserved architecture (score 2; Fig. 5E). The reduction was significant versus control (ANOVA P = 0.0032; combination versus control, Tukey P = 0.0021), as was GSK212 monotherapy (Tukey P = 0.031), though the combination did not significantly exceed either monotherapy at this sample size (non-parametric confirmation in table S8). Combination treatment therefore limits early structural injury during severe ambient storage, with a consistent but non-significant reduction in biochemical markers.

To confirm the dose, we performed dose-ranging under extended hypothermic storage for 18 hours at 12°C or 4°C. At 12°C we compared 0.3×, 1×, and 3× against untreated control (1× = 2 µM GSK212 and 3 µM AP39 in the perfusate, with a 600 nM AP39 bolus at reperfusion; all components scaled together); at 4°C we compared 1× and 3× against control. At both temperatures 1× gave the lowest transaminases and 3× performed no better (fig. S9, A and B; fig. S10, B and C). At 4°C (n = 3), 1× yielded significantly lower AST and ALT than 3× (one-way ANOVA with Tukey post hoc; AST P = 0.030, ALT P = 0.040; table S9, A and B). The response was biphasic about 1×: transaminases fell below control at 1× (AST 1888 versus 3101 U/L; ALT 1764 versus 2603 U/L) and rose above it at 3× (AST 3583 U/L; ALT 3407 U/L), though neither comparison with control reached significance at this sample size. This loss of protection on escalation matches the known concentration-dependent behavior of H₂S donors, which shift from cytoprotective to inhibitory(*26*). We adopted 1× for all subsequent transplantation experiments. The 12°C cohort was exploratory, with one to three animals per group, precluding statistical testing. Its histology followed the same rank order, with 1× yielding the lowest Suzuki score (fig. S9, C and D), but because that cohort was stored colder and for more than three times the duration of the room-temperature experiments, its smaller effect size reflects a different and more severe challenge rather than a discrepant result.

To test functional viability, we monitored a separate parallel survival cohort (**5 h RT** preservation) over 14 days (Fig. 5F; n = 4 per group), with daily welfare checks and deaths assigned to the preceding check interval. Control and AP39 grafts failed rapidly, with all recipients found deceased at the first check (0.5 days). GSK212 and the combination each brought 25% of recipients to day 14. Among non-survivors, combination-treated animals died later (1.5 to 9.5 days) than GSK212-alone animals (0.5 to 1.5 days), and the combination cohort held 50% survival through day 9 versus 25% for GSK212. A Cox proportional-hazards model incorporating GSK212, AP39, and their interaction (likelihood-ratio χ²(3) = 10.23, P = 0.017; table S10A) confirmed that GSK212 significantly reduced mortality hazard (HR 0.13, 95% CI 0.02 to 0.87, P = 0.04), while AP39 had no significant effect (P = 0.72) and no GSK212×AP39 interaction was detected (P = 0.70; table S10B). The hazard ratio corresponds to an approximately eightfold reduction in mortality risk attributable to GSK212, though the wide confidence interval reflects the limited cohort size.

Transaminases in surviving recipients at day 14 confirmed terminal graft recovery: the single survivor in each of the GSK212 and combination cohorts normalized biochemically, with AST of 91 and 84 U/L (Fig. 5G) and ALT of 68 and 64 U/L (Fig. 5H). Because a single recipient survived in each arm, these values establish that full biochemical recovery is attainable rather than quantifying how often it occurs.

## Discussion

Decoupling tissue viability from continuous perfusion or hypothermia remains a central challenge in transplantation. Static cold storage slows metabolic kinetics but mitigates IRI only briefly(*3–7*), while NMP maintains physiological parameters *ex vivo* at severe logistical and financial cost(*8*). Neither alters the tissue’s intrinsic metabolic drive. True organ biostasis requires shifting from environmental manipulation to cell-autonomous control.

Prior attempts at synthetic hypometabolism have remained tethered to the gas phase. H₂S gas induced an apparent suspended-animation state in mice(*17*), but the effect tracked the reduced ambient oxygen (∼17.5% O₂) rather than any direct pharmacological action, and later mammalian models showed limited efficacy attributable to hypoxia-induced anapyrexia(*18, 41–45*). Classical mTOR-mediated diapause has been demonstrated only under 5% O₂(*19, 20*), and our data confirm that INK128 arrested embryos at 5% O₂ but was lethal at 21% (Figs. 3 and 4, and fig. S3).

Mammalian embryonic diapause nonetheless offers a conserved blueprint for reversible arrest(*12, 14, 15*), shown directly by cross-species transfer: non-diapausing sheep embryos arrest within diapausing murine hosts and resume development on transfer to receptive sheep uteri (*46*). Pharmacological translation has been limited by target tractability and by the hypoxia requirement. Myc depletion induces a reversible diapause-like state in mouse blastocysts for <24 h(*47*), but Myc lacks direct inhibitors(*48*) and available tools act indirectly on Myc-Max dimerization at micromolar concentrations. We reasoned that hypoxia substitutes for a missing second block: mTOR inhibition alone leaves PI3K free to reactivate AKT and restore proliferation (*21, 22*), whereas blocking both nodes should maintain arrest at ambient oxygen. Our earlier finding that TGF-β drives murine HSCs into a reversible hibernation-like state at 21% O₂ through PI3K-AKT suppression and FoxO3a accumulation supported this logic(*23*).

The screen bore this out. Combined pan-PI3K and mTOR inhibition suppressed proliferation synergistically while isoform-selective PI3K inhibition was merely additive, implicating parallel PI3K signaling as the compensatory route permitting proliferation under mTOR blockade alone, and explaining why single-node approaches require hypoxia(*22, 49, 50*). Among dual-node agents, GSK212(*24, 25*) uniquely combined sub-nanomolar inhibition of all four class I PI3K isoforms and both mTOR complexes with a usable reversibility window, recovering even from near-maximal suppression at 500 to 1000 nM where every comparator failed. RapaLink-1 illustrates the opposite failure mode: its bivalent design confers durable engagement, since binding to abundant intracellular FKBP12 sustains inhibition after washout(*51*), making it unsuitable where reversibility is the objective. Dual-node inhibition is therefore a viable route to biostasis independent of oxygen control. To counter acute surgical ischemia we paired this metabolic brake with AP39, which delivers H₂S directly to mitochondria to reversibly inhibit Complex IV(*26–28*) as a mitochondrial protectant.

Combining the two yielded TorPause. Transcriptional profiling of stored organs supported a torpor-like state, with one qualification: storage alone suppressed interferon programs in all four organs, and GSK212 and the combination deepened that suppression while additionally suppressing the allograft-rejection signature, which was unchanged or elevated in controls (figs. S5 to S7). The immune quiescence of natural hibernation is governed by a profound drop in body temperature toward ambient values(*52*); TorPause reproduced this signature at 23°C, without the deep cooling hibernators require and without refrigeration. Because these signatures were measured before reperfusion in a non-transplant preparation, they indicate pre-conditioning toward a lower-inflammatory state rather than an altered rejection outcome.

We chose our RT model for two reasons. It isolates the source of metabolic suppression, since at ambient oxygen any hypometabolism must arise pharmacologically or from the modest cooling of room temperature, not from the hypoxia on which earlier H₂S models depended. And it targets a clinical blind spot: neither cold storage nor NMP protects the graft during surgical anastomosis. In the RT cohort that blind spot spans the whole experiment rather than a brief window at its end, since after the initial flush the graft is held in static storage at RT for 5 h and through implantation to reperfusion. Under this challenge TorPause significantly reduced structural liver injury, with GSK212 the significant driver of improved recipient survival. Because the intervention is a single flush into solution already in clinical use, adoption requires no capital equipment, no perfusion expertise, and no change to organ-handling workflow, the barriers that have limited machine perfusion to well-resourced centers. Acting cell-autonomously rather than by altering the storage environment, TorPause should also be compatible with machine perfusion, though whether it adds to perfusion remains untested. The combination gave the most favorable histology and survival but did not significantly exceed either monotherapy at this sample size, and the reduction in transaminases was consistent but not significant. TorPause therefore mitigates rather than abolishes IRI at the durations and temperatures tested; among recipients surviving the acute window, biochemistry normalized fully by day 14. One design limitation is that grafts remained in drug-containing solution throughout storage, so retained intracellular drug cannot be separated from continued extracellular exposure. A flush-wash-store protocol would isolate the cell-autonomous component and better mirror clinical practice; this is the immediate next step.

AP39 is already established as protective in organ preservation, but under conditions that differ fundamentally from the challenge addressed here. It improves outcomes in cold static storage of kidney (*39*) and heart(*38*), and during normothermic kidney perfusion(*53*). In each setting a second factor was already limiting injury: hypothermia suppresses metabolic demand during cold storage, while perfusion actively supplies oxygen and substrate. Several of these studies also used *ex vivo* perfusion as a surrogate rather than orthotopic implantation with survival endpoints. AP39 was therefore never tested as the sole source of protection in a warm graft held in static storage. Here AP39 alone was insufficient, conferring structural protection only when paired with GSK212, suggesting that mitochondrial H₂S delivery and metabolic suppression address complementary, non-redundant components of warm IRI. Consistent with the biphasic activity of H₂S donors(*26*), escalating beyond 1× diminished protection: the 3× dose performed significantly worse than 1× at 4°C and exceeded untreated control, identifying 1× as an optimum rather than a lower bound. Because 1× was optimal at 4°C and 12°C as well as at RT, the formulation required no retuning across conditions ranging from conventional 18-hour cold storage to 5 hours at RT.

TorPause requires no perfusion circuit and integrates into existing static storage protocols as a simple pharmacological flush, a profile that could broaden access to advanced preservation. Uncoupling metabolic suppression from environmental oxygen control provides a versatile platform, and the same torpor-like signature across all four perfused organs offers preliminary support that this engagement is not liver-restricted, though functional confirmation in each organ will be required. Extending this pause to systemic applications, including whole-body perfusion for simultaneous multi-organ procurement, presents a plausible trajectory for treating trauma, stroke, and acute ischemic injury.

## Material and Methods

### Animals

All animal experiments were performed in accordance with the ethical guidelines and standards set by the Stanford University Administrative Panel on Laboratory Animal Care, which reviewed and approved the protocols (#29042 and #9821) related to animal handling and care. Animals were housed under specific pathogen-free conditions in a temperature- and humidity-controlled environment with a 12-hour light-dark cycle, with unrestricted access to water and standard chow.

Male CD-1 mice (15-18 weeks of age, 50-55 g) used for whole-organ perfusion, and CD-1 mice used for embryo production, were sourced from Charles River Laboratories, Wilmington, MA. For embryo collection, CD-1 females (3-4 weeks) were superovulated and mated with CD-1 stud males, and embryos were recovered as described below.

Male Lewis rats (7-8 weeks of age) used for orthotopic liver transplantation were sourced from Charles River Laboratories, Wilmington, MA.

### Embryo Culture

Mouse embryos were prepared following the protocol described(*54, 55*). Briefly, embryos were collected from superovulated CD-1 females (3-4 weeks) paired with CD-1 stud males. Successful mating was confirmed by checking for the presence of a vaginal plug on embryonic day or day post-coitum 0.5 (DPC 0.5). Embryos were collected at the 2-cell stage (DPC 1.5) by flushing out the oviducts and rinsed three times with M2 medium to remove residual fluid. 2-cell stage embryos were transferred directly to fresh KSOM-AA medium without phenol red (reagents sourced from CytoSpring, Mountain View, CA; Cosmo Bio, Tokyo, Japan; or MilliporeSigma, Burlington, MA across experimental batches) for imaging (37°C, 21%O₂, and 5% CO₂.). Embryos were cultured in non-tissue culture treated 96-well U-bottom plates (351177, Corning Incorporated, Corning, NY), with one embryo per well and the outer wells filled with Milli-Q water to prevent edge effects. KSOM-AA without phenol red was centrifuged immediately before use to pellet particulate debris, and 50 µl of the clarified medium was added to each well. All empty wells were filled with 250 µl of deionized water and the plate was equilibrated for at least 1 h at 37 °C and 5% CO₂ before embryos were loaded.

For extended culture experiments (Fig. 4), embryos were generated by *in vitro* fertilization using CARD MEDIUM per manufacturer’s protocol (KYD-003-EX, Cosmo Bio, Tokyo, Japan) to ensure controlled developmental timing, and cultured to the morula stage (DPC 2.5) prior to plating for imaging.

### Embryo live imaging

Live imaging of embryos was performed on an Operetta High Content Screening System (PerkinElmer, Waltham, MA) fitted with a live-imaging chamber maintained at 37 °C, 5% CO₂, 21% O₂, at 30-min intervals for the dose-titration experiment (1.5 to 6.5 DPC) and 3-h intervals for the extended pausing experiment (2.5 to 11 DPC). All imaging data were processed and analyzed using FIJI(*56*) as outlined below.

Images were acquired with a 10×/0.3 NA objective in non-confocal mode, at a single field and a single z-plane positioned 45 µm above the well bottom. Each image is 1360 × 1024 pixels at 0.993 µm per pixel, giving a 1.35 × 1.02 mm field of view. Depending on the experiment, images acquisition intervals varied (15-min to every 3-h). Only the transmitted-light brightfield channel was used in this study; the system acquires epifluorescence channels in parallel, so fluorescent reporters could be imaged alongside brightfield in future experiments without altering the acquisition or analysis described here.

A 10× objective was chosen over higher magnification for two reasons. First, embryos do not settle reproducibly at the center of a U-bottom well, and the well is not re-centered between timepoints; in a representative experiment, just four embryos occupied a region spanning approximately 400 × 350 µm across wells. The 1.35 × 1.02 mm field of a 10× objective therefore captures the embryo wherever it comes to rest, whereas a 20× objective would halve the field in each dimension and risk excluding embryos that settle off-center. Second, because only a single z-plane was acquired per timepoint, the depth of field of the objective determines how much of the embryo remains in acceptable focus. This matters most after blastulation, when the embryo expands and thickens: with the greater depth of field of the lower-magnification objective, the majority of the blastocyst remains in focus in the single acquired plane, and the outer boundary stays sharp enough to segment.

### Embryo image processing

Analysis was performed with FIJI(*56*) using macros written to organize, segment, measure, and crop the image series (fig. S2A). The brightfield channel was used for all segmentation and tracking.

For each well and timepoint, the raw brightfield image was thresholded using the MaxEntropy auto-threshold method and converted to a binary mask. The mask was then processed through a fixed sequence of morphological operations: a short dilate-fill-erode cycle to close the granular interior of the embryo into a single solid object, followed by 10 rounds of erosion and 12 rounds of dilation (fig. S2A).

The erosion step is essential. Wells contain debris and the plate surface carries scratches, both of which threshold at similar intensity to the embryo boundary and appear as elongated or punctate objects in the raw mask (white arrows, top right, fig. S2A). Because these features are thin, successive erosion removes them entirely while the embryo, being compact, survives; the subsequent dilation restores the embryo to approximately its original size. Residual objects were excluded by a particle-size filter (minimum approximately 4,300 px², equivalent to a circular object of roughly 74 µm diameter), leaving a single mask per well whose outline corresponds closely to the embryo (blue line, fig. S2A).

This mask serves three purposes. First, it defines the region over which morphological measurements are made for each embryo at each timepoint: size, taken as the mask area, and roundness, for which 1 is a perfect circle (fig. S2B). Second, it provides a region within which fluorescence could be quantified on a per-embryo basis if needed in future experiments. Third, it yields the embryo’s position at every timepoint, which allows the embryo to be tracked over the time course and the crop window to be placed on it (green box, fig. S2A).

Tracking proceeds as follows. Where filtering leaves more than one candidate mask in a well, a single object must be chosen: at the first timepoint the roundest candidate is taken as the embryo, and at every subsequent timepoint the candidate whose centroid lies nearest the previous position is taken, so the identity of the tracked object is maintained even when additional objects are present. Candidates larger than roughly half the field of view are disregarded at every timepoint, since an object of that size reflects a failed threshold rather than an embryo. The resulting position trace is smoothed with a five-point running median (three-point at the second and penultimate timepoints) to suppress frame-to-frame jitter, and the smoothed coordinates define the center of a fixed-size square crop applied at that timepoint.

Adaptive cropping is required for two reasons. First, embryos do not begin in the same place: at 2.5 DPC their positions already differ by several hundred micrometers between wells (fig. S2C). Second, embryos move within the well over the time course, particularly after hatching from the zona pellucida, from approximately 3.75 DPC onward (fig. S2B and fig. S2C). A single fixed crop window, applied to every well and every timepoint, must therefore be large enough to contain every embryo at every position it occupies - a 410 × 410 px (407 × 407 µm) window for the experiment shown, in yellow in fig. S2D. Cropping each well and timepoint independently, re-centered on the tracked position, instead requires only a 250 × 250 px (248 × 248 µm) window (green, fig. S2D), a 2.7-fold reduction in area. The adaptive crop therefore yields more closely framed images in which each embryo remains centered throughout the time course, which makes morphological changes easier to follow across the series and in the resulting time-lapse videos.

### mESC Derivation and Culture

For mESC derivation, naturally mated CD-1 female mice (3-4 weeks) with confirmed vaginal plugs at DPC 0.5 were allowed to develop until DPC 3.5. Blastocyst-stage embryos were directly flushed out and rinsed in M2 medium. Single embryos were transferred into 12-well plates (Corning Incorporated, Corning, NY) onto mitomycin-c treated feeder layers using N2B27-2iL medium as previously described(*29, 30*).

The mESC line SUN107’s derivative (SUN107.4) was employed for small molecule screening and titration experiments. Specifically, the SUN107.4 line, a fluorescently labeled single-cell cloned mESC line derived from SUN107, was utilized to ensure consistent mean fluorescent intensity (MFI). This line was tagged with a PiggyBac vector carrying tdTomato, and a clone exhibiting optimal fluorescence and developmental potential was selected following verification through mouse-mouse chimera assays (data not shown). Cells were cultured and maintained on feeder layers in N2B27-2iL medium, passaged at a density of 7,000-10,000 cells/cm² every 3-4 days to preserve growth and pluripotency.

SGE2 is a male, germline-competent mESC line derived in our laboratory from C57BL/6 transgenic mice constitutively expressing EGFP, and is competent to contribute to interspecies chimeras(*57*). SGE2’s were maintained and passaged as described above at 37 °C and 5% CO₂ on mitomycin C-treated MEF feeder layers in GMEM supplemented with 0.5× GlutaMAX, 1× non-essential amino acids, 1× sodium pyruvate, 50 µM β-mercaptoethanol (all from Thermo Fisher Scientific, Waltham, MA), 10% fetal bovine serum (Sigma-Aldrich, St. Louis, MO), human LIF (20 ng/ml, PeproTech (Thermo Fisher Scientific), Waltham, MA), and 2i (1 µM PD0325901, 3 µM CHIR99021, MedChemExpress, Monmouth Junction, NJ) at final concentration.

### Small Molecule Screening and Dosage Titration

All small molecules used for screening, as listed in Supplementary table S1, were purchased from SelleckChem and pre-dissolved at a concentration of 10 mM in dimethyl sulfoxide (DMSO) as described previously(*58*). For screening assays, SUN107.4 mESCs were seeded at a density of 10,000 cells/cm² in N2B27-2iL medium, with each inhibitor added at a final concentration of 1 μM, prepared by diluting the stock solution in N2B27-2iL medium, ensuring the final DMSO concentration did not exceed 0.1%. Cells were imaged at T1 [12h post-seeding] and subsequently every 24 hours. Following the initial screening period, cells were washed three times with Dulbecco’s phosphate-buffered saline (dPBS; Thermo Fisher Scientific, Waltham, MA) to remove residual inhibitors and returned to fresh N2B27-2iL medium, with imaging continuing at 24-hour intervals as described in Results.

For dosage titration experiments, small molecules were sourced from MedChemExpress (MedChemExpress, Monmouth Junction, NJ). All lyophilized powders were dissolved in anhydrous DMSO (Thermo Fisher Scientific, Waltham, MA) and stored as single-use aliquots at concentrations exceeding 10,000X stock at -80°C until use. SUN107.4 mESCs were seeded at a density of 10,000 cells/cm² in N2B27-2iL medium supplemented with the indicated concentrations of small molecules, ensuring the final DMSO concentration did not exceed 0.1%. Cells were imaged at T1 and subsequently every 24 hours over a 5-day exposure period. Cells were then washed three times with dPBS to remove residual inhibitors and returned to fresh N2B27-2iL medium, with imaging continuing at 24-hour intervals for an additional 5 days to monitor proliferation recovery.

### Whole-organ perfusion and static storage

CD-1 mice were anesthetized with 3% isoflurane and systemically heparinized 1 IU/g (Fresenius Kabi, Lake Zurich, IL). Organs were flushed *in situ* via abdominal aorta with 10 mL of room temperature (23 ± 2°C, RT) University of Wisconsin (UW) solution; warm ischemia time (WIT) at procurement was approximately 5-6 min for all animals. Storage duration is defined as the interval from interruption of donor hepatic blood flow to removal of the graft from storage for implantation. This interval encompasses procurement, flush, and static storage. Perfusate defined the treatment group: UW alone (Control), or UW containing AP39 (3 µM, MedChemExpress, Monmouth Junction, NJ), GSK212 (2 µM, MedChemExpress, Monmouth Junction, NJ), or AP39 (3 µM) plus GSK212 (2 µM) (Combination); small molecules were diluted from DMSO stocks, with Control perfusate matched for DMSO (0.1% v/v). Heart, lung, liver, and kidney were then excised, submerged in 20 mL of the identical group-matched solution (30 mL total per animal), and stored for 5 h at RT. A separate baseline group (Sham) was perfused and snap-frozen immediately without storage. At the end of storage, and before any reperfusion, tissue from each organ was minced, snap-frozen on dry-ice, and stored at -80 °C. Three biological replicates (independent animals) were collected per organ and condition, giving 60 samples (4 organs × 5 conditions [Sham, Control, AP39, GSK212, Combination] × 3 replicates).

### Mouse whole organ transcriptomics and bio-informatics pipeline

RNA extraction, library preparation, and sequencing were performed by Novogene (Novogene, Sacramento, CA). Briefly, strand-specific poly(A) libraries (NEBNext Ultra II Directional) were sequenced on an Illumina NovaSeq X Plus (150 bp paired-end) to a mean depth of 57.8 million raw read pairs per sample (range 42.7–80.7 M; ≥6 Gb/sample). Sequencing quality was high across all samples (mean Q30 95.9%, Q20 99.0%, error rate 0.01%, GC 47.3%).

#### Read processing and alignment

RNA-seq reads were processed and aligned as described previously(*59*), using the current versions of the respective software packages. In brief, reads were quality- and adapter-trimmed with Trim Galore (Cutadapt/FastQC) and screened for contamination with FastQ-Screen (Bowtie2). Trimmed reads were aligned with STAR v2.7.4a to the mouse GRCm39 primary assembly indexed against GENCODE vM37 annotation (--sjdbOverhang 100), using --outFilterType BySJout, --outFilterIntronMotifs RemoveNoncanonicalUnannotated, and --outSAMstrandField intronMotif. Alignments were sorted and indexed with SAMtools v1.17 and assessed with Qualimap v2.2.2d (strand-specific-reverse, paired-end). Gene-level counts were generated with HTSeq-count v0.11.2 (--stranded reverse, --additional-attr gene_name) against the same GENCODE vM37 annotation, and per-sample QC aggregated with MultiQC v1.29.

#### Differential expression and gene-set enrichment

Count-level analysis used DESeq2 (implemented in PyDESeq2 v[0.5](*60*)). Genes with CPM > 1 in at least three samples were retained (21,294 genes) and duplicate gene symbols collapsed by summation. Differential expression was computed within each organ, contrasting each stored condition against the fresh no-storage Sham baseline (Wald test, Benjamini-Hochberg FDR). Counts were variance-stabilized (VST) for ordination; principal-component analysis on the 2,000 most variable genes confirmed that samples separated primarily by organ of origin. Gene-set enrichment analysis(*61, 62*) was performed in preranked mode (gseapy)(*63*) on genes ranked by the DESeq2 Wald statistic, against the MSigDB Mouse Hallmark collection(*64*) (mh.all.v2024.1.Mm; 50 gene sets, 1,000 permutations), with significance at FDR q < 0.05. Negative normalized enrichment scores denote suppression of a program relative to the healthy Sham baseline. The normalized expression matrix and full enrichment statistics are provided in table S7.

### Rat orthotopic liver transplantation

The protocol for orthotopic liver transplantation was adapted from established methods(*65*). Briefly, after donor rats were heparinized with 1 IU/g (Fresenius Kabi, Lake Zurich, IL), the descending thoracic aorta was clamped, and the supra-hepatic inferior vena cava (SHIVC) was cut. The liver was flushed using a catheter inserted into the abdominal aorta with 50 mL of phosphate-buffered saline (PBS; Thermo Fisher Scientific, Waltham, MA), followed by 20 mL of University of Wisconsin (UW) solution (Bridge to Life Ltd., Northbrook, IL). The liver was then excised and immediately immersed in a basin of UW solution. For the treatment group, 2 µM of GSK212 and 3 µM of AP39 were added to the UW solution. A cuff fashioned from a 14-gauge ethylene tetrafluoroethylene (ETFE) catheter was applied to the portal vein (PV) during back-table preparation. The liver was stored in UW solution under one of three conditions: 5–6 hours at room temperature (23 ± 2°C), 18 hours at 12°C, or 18 hours at 4°C.

Immediately before transplantation, the liver was flushed with 5 mL of PBS. For the 18-hour storage cohorts (12°C and 4°C) only, dual-treated (GSK212+AP39) livers received an additional 600 nM AP39 supplement in this pre-unclamping PBS flush, a dosage twice that used in previous preservation studies, to assess whether reperfusion-timed dosing conferred additional protection based on the cardioprotective effects of slow-releasing H_2_S donors observed in prior research(*66*). This supplemental bolus was not administered in the 5-hour RT cohort, and provided no significant additive protective effect relative to combination treatment during storage alone (see Results, fig. S9-10).

The liver was then orthotopically transplanted into the recipient, with the SHIVC anastomosed end-to-end using continuous 7-0 polypropylene sutures. The PV was reconstructed using a cuff technique, and reperfusion was achieved within 16 min of clamping, defining an anhepatic phase of ≤16 min during which the graft was at ambient temperature and unperfused. Total warm ischemia therefore comprised the storage interval plus this implantation window. The infra-hepatic IVC (IHIVC) was anastomosed end-to-end with continuous 8-0 polypropylene sutures, and the hepatic artery (HA) was reconstructed by inserting the recipient’s proper HA into the celiac artery of the liver graft. The bile duct was reconstructed end-to-end using a 24-gauge ETFE catheter as a stent.

### Sampling and Biochemical Analysis for liver transplantation

At 2 hours post-transplantation, serum and liver samples were collected. Serum levels of AST and ALT were analyzed at the Animal Diagnostic Laboratory at Stanford University. Liver tissue samples were fixed in 4% formaldehyde for 2 days at room temperature, followed by 70% ethanol before being shipped to Histo-Tec Laboratory, Hayward, CA for further processing. Briefly, samples were embedded in paraffin and sectioned at 5 µm for H&E staining. Histological assessment of IRI injury, including necrosis, sinusoidal congestion, and centrilobular ballooning, was performed using the modified Suzuki score(*67*).

### Statistical Method

Synergy in the combinatorial screen was assessed with the Bliss independence model(*68*). For each pair, the expected combined inhibition was calculated as E = A + B − (A × B), where A and B are the fractional inhibitions of each single agent relative to untreated control, and the Bliss score was taken as the difference between observed and expected inhibition in percentage points, with positive values indicating synergy and values near zero indicating additivity. Inhibition was derived from confluence at T4, and all 72 pairwise combinations were scored (table S2, A and B). Because the classical Bliss score does not account for response variability and can therefore overcall synergy(*69*), we did not interpret scores below the replicate variability of the assay; variability among untreated control wells was approximately 7.5 percentage points, so scores within that range were treated as additive. Statistical analyses were performed in Python (v3.12.3) using scipy (v1.17.1) for one-way ANOVA and the Kruskal-Wallis test; statsmodels (v0.14.6) for Tukey’s HSD post hoc testing and Holm-Bonferroni correction; scikit-posthocs (v0.14.0) for Dunn’s post hoc test; and lifelines (v0.30.3) for Kaplan-Meier estimation, log-rank testing, and Cox proportional hazards regression. For the embryo survival experiments (Fig. 3, D and E, and Fig. 4B), Kaplan-Meier estimation, omnibus log-rank (Mantel-Cox) testing, and pairwise log-rank comparisons with Holm-Bonferroni correction were performed in lifelines; pairwise results are reported in tables S4A and S5A. GraphPad Prism (v11.0.2) was used for graph preparation. RNA-seq differential expression and gene-set enrichment were analyzed with DESeq2 (PyDESeq2) and preranked GSEA (gseapy) as detailed above, with significance controlled by the Benjamini-Hochberg FDR. Data are presented as mean ± SD unless otherwise indicated, and P < 0.05 was considered statistically significant (with RNA-seq enrichment thresholded at FDR q < 0.05). Exact sample sizes are stated in each figure legend.

---

**Supplementary table S1.** List of small molecules used for the screen, with target, pharmacological class, potency, mechanism, and clinical status.

**Supplementary table S2.** Cell confluence across the combinatorial inhibitor screen (A) and Bliss synergy analysis (B) (related to Fig. 1C and fig. S1).

**Supplementary table S3.** Concentration-response regression (A) and post-washout recovery regression (B) for the mESC dose titration (related to Fig. 2).

**Supplementary table S4.** Survival statistics (A) and imaging timeline (B) for GSK212-paused embryos under normoxia (related to Fig. 3, D and E).

**Supplementary table S5.** Survival statistics (A) and imaging timeline (B) for extended embryo pausing (related to Fig. 4B).

**Supplementary table S6.** Raw viability and live-cell counts for sub-normothermic mESC storage (related to fig. S4).

**Supplementary table S7.** RNA-seq processed data for the organ transcriptomic analysis (related to figs. S5 to S7).

**Supplementary table S8.** Hepatocellular injury following 5-hour ambient storage (related to Fig. 5, B, C, and E).

**Supplementary table S9.** Dose-ranging of combination treatment following 18-hour storage at 4°C: omnibus (A) and pairwise (B) tests (related to fig. S10).

**Supplementary table S10.** Cox proportional-hazards analysis of recipient survival: model summary (A) and per-term hazard ratios (B) (related to Fig. 5F).

**Movie S1.** Automated embryo segmentation and tracking. Time-lapse acquisition showing the image-processing workflow applied at each timepoint: brightfield images are thresholded to a binary mask, filtered by erosion and dilation to remove debris and plate scratches, and the resulting mask used to locate each embryo and record its shape, size, and position over time (related to fig. S2, A to C).

**Movie S2.** Adaptive cropping keeps each embryo centered. Side-by-side comparison of uncropped acquisition, a single fixed crop window sized to retain all embryos at every timepoint (yellow box), and adaptive cropping in which each well and timepoint is cropped independently (green box), showing that adaptive cropping produces more closely framed images in which each embryo remains centered despite drift during culture (related to fig. S2D).

**Movie S3.** Time-lapse imaging of embryo pausing under normoxia, from which the micrographs and Kaplan-Meier data in Fig. 3, C to E, were derived. Each panel follows the same individual embryo in a fixed well position throughout; embryos were not pooled, replaced, or resampled between timepoints. Embryos were imaged at 30-min intervals, and viability was scored at 24-h intervals from the complete time-lapse series.

**Movie S4.** Time-lapse imaging of extended embryo pausing, from which the micrographs and Kaplan-Meier data in Fig. 4, A and B, were derived. Each panel follows the same individual embryo in a fixed well position throughout. Embryos were imaged at 3-h intervals, and viability was scored at 24-h intervals from the complete time-lapse series. Each acquired frame is displayed for approximately 0.5 seconds; frames were duplicated without interpolation for playback, and no time points were interpolated or omitted.

## Author Contributions

JB conceived and designed the study, performed experiments, analyzed data, and wrote the manuscript with input from all authors. JB and TT performed the whole-organ perfusion experiments, and JB performed the RNA-seq analysis. TT optimized and conducted liver transplantations, analyzed data, and interpreted results. FS developed the imaging platform used for small-molecule screening in mESCs and embryos, and conducted embryo culture and mESC derivation. TN assisted with embryo experiments. HM independently validated identified small molecules in mESCs and embryos. JZ performed mESC experiments. MR helped optimize initial mouse perfusion experiments. CTC and TN contributed to the discussion of mTORi in whole organ pausing. COE and HN supervised the study and provided guidance on the overall project. All authors read and approved the final manuscript.

## Acknowledgements

We thank K. C. Chan and S. Homma for laboratory and administrative support, M. Rodriguez for assistance with small-molecule titration experiments, and all members of our laboratories for insightful discussions. Figures were created in part using BioRender (biorender.com). AI-assisted language editing was used in preparing the manuscript. This work was supported by grants from the Dr. Ralph and Marian Falk Medical Research Trust – Catalyst Awards Program (2023) to JB, COE, and HN. JB was supported in part by an international postdoctoral grant (2017-0034) from the Swedish Research Council and the Assar Gabrielsson Foundation, Sweden. TT was supported by the Stanford Transplant and Tissue Engineering Center of Excellence fellowship and the Overseas Research Fellowships of the Japan Society for the Promotion of Science (JSPS). FS was in part supported by the National Science Foundation Graduate Research Fellowship; and the Pat Tillman Fellowship. This work was partly supported by the Joint Research Program of the Joint Usage/Research Center at the Institute of Development, Aging and Cancer, Tohoku University to HM. TN was supported by the Japan Society for the Promotion of Science (JP18K14602 and JP18J00499). The work was supported in part by the Arnold & Barbara Silverman Endowment at the Stanford University School of Medicine, and by the Nancy & Richard Robbins Family Foundation to COE. Part of this project was supported by CIRM (DISC2P-11662), NIH (R01DK121851) and grants from the Japan Agency for Medical Research and Development (AMED) under Grant Numbers JP22bm1004002 and JP23bm1123041h0001 to HN.

## Data and materials availability

Numerical source data underlying all main and supplementary figure panels are provided in supplementary tables S1 to S10. RNA-sequencing raw data will be deposited in the Gene Expression Omnibus upon publication. All other data are available in the main text or the supplementary materials.

## Supplementary Materials

Materials and Methods

Figs. S1 to S10

Tables S1 to S10

Movies S1 to S4

