## Supplemental Figures for "TorPause: Efficient Static Organ Preservation for Transplants"

**Fig. S1**

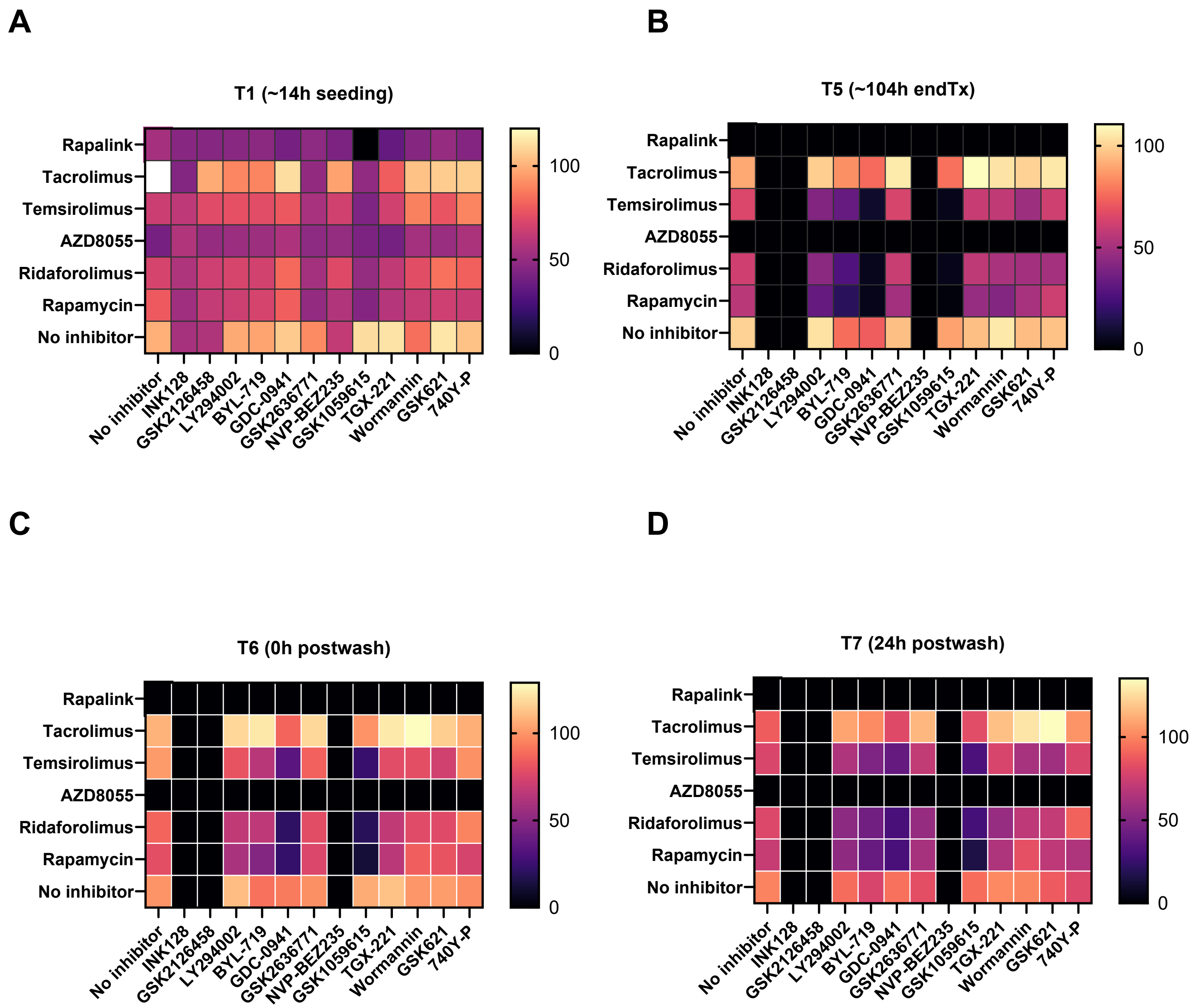


**Fig. S1. Time-resolved combinatorial inhibitor screen showing pausing and washout behavior.** Related to Fig. 1. Mouse ESCs were seeded with single inhibitors (1 µM) or pairwise combinations (1 µM of each agent) in a matrix of six mTOR-pathway inhibitors (rows) against twelve PI3K- and related-pathway inhibitors (columns), and imaged longitudinally. Heat maps show cell confluence (colony number × colony size) as a percentage of the untreated control (mean of four control wells) at four timepoints: (**A**) T1 (~14 h post-seeding, baseline), (**B**) T5 (~104 h, end of treatment), (**C**) T6 (~0 h after inhibitor washout), and (**D**) T7 (~24 h after washout). The T4 timepoint (~90 h) is shown in Fig. 1C. In each panel, the top row and left column show single-agent (No inhibitor) references and the interior shows combinations. At seeding (T1), confluence was comparable across conditions, confirming equal plating; by the end of treatment (T5), potent single agents and combinations had suppressed confluence to near-baseline. Following washout, moderately suppressed cultures resumed proliferation (T6–T7), whereas the most deeply suppressed colonies did not recover, indicating that the deepest pharmacological arrest under normoxia resulted in cell loss rather than a reversible pause. RapaLink-1's failure to recover reflects its bivalent design rather than incomplete washout, since binding to abundant intracellular FKBP12 sustains target engagement after drug removal(*1*). Raw data for confluence and percent-of-control values for all timepoints and conditions are provided in table S2A.

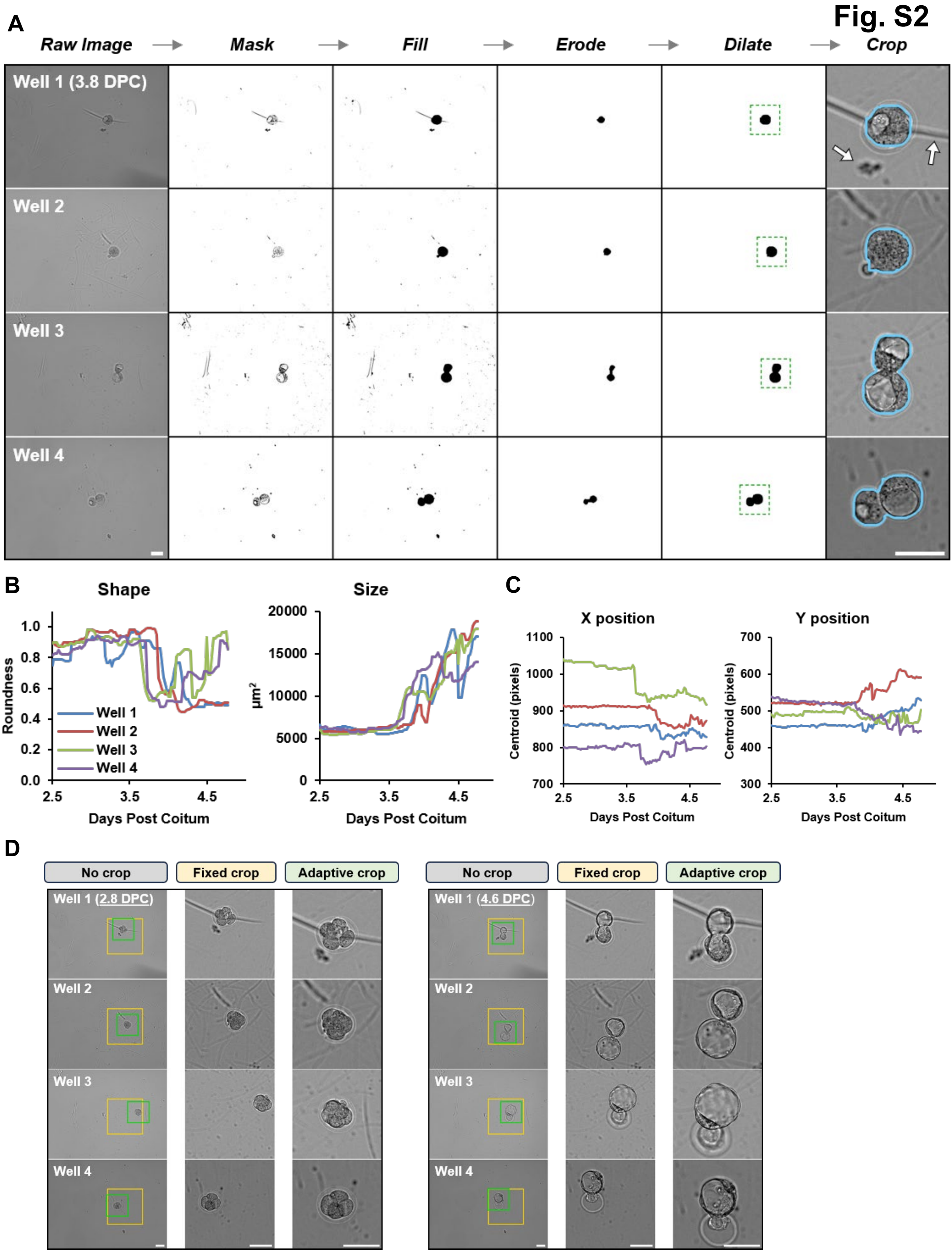

Fig. S2
A
Raw Image
Mask
Fill
Erode
Dilate
Crop
Well 1 (3.8 DPC)
Well 2
Well 3
Well 4
B
Shape
Roundness
Well 1
Well 2
Well 3
Well 4
Days Post Coitum
Size
µm²
Days Post Coitum
C
X position
Centroid (pixels)
Days Post Coitum
Y position
Centroid (pixels)
Days Post Coitum
D
No crop
Fixed crop
Adaptive crop
Well 1 (2.8 DPC)
Well 2
Well 3
Well 4
No crop
Fixed crop
Adaptive crop
Well 1 (4.6 DPC)
Well 2
Well 3
Well 4

### Fig. S2

**Fig. S2. Embryo imaging and analysis pipeline. (A)** Image processing workflow. Brightfield images are converted to a binary mask, which is then filtered by a series of erode and dilate commands: erosion removes thin objects such as plate scratches and debris (white arrows), and dilation restores the embryo to its original size. The resulting mask locates the embryo at each timepoint and defines the crop window (green box). The mask outline (blue) is shown overlaid on the cropped image (right column). **(B)** Embryo morphology over time, quantified from the mask as roundness (1 = perfect circle) and size (mask area). Each trace is one embryo (n = 4). **(C)** Position of the same four embryos over time. Embryos begin at different positions and move during culture. Coordinates are given in pixels of the raw image, 1360 (X) × 1024 (Y), corresponding to 1.35 × 1.02 mm. Traces are colored as in B. **(D)** Adaptive cropping produces more closely framed images in which each embryo remains centered. The yellow box shows the smallest single window that keeps all four embryos in frame at every timepoint (410 × 410 px). The green box shows adaptive cropping, in which each well and timepoint is cropped independently (250 × 250 px). Scale bars, 100 µm.

### Fig. S3

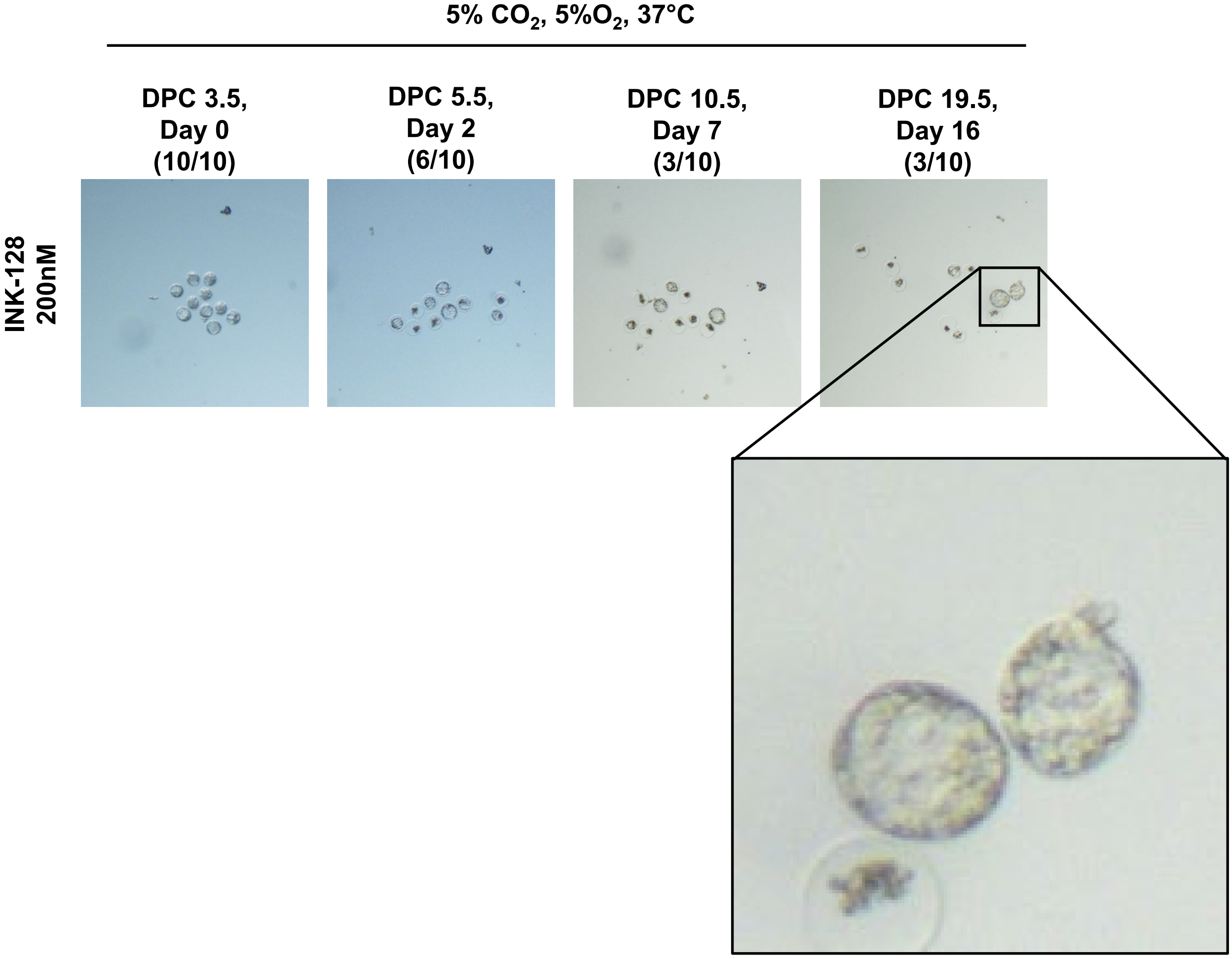


**Fig S3: INK128 sustains embryonic pausing under hypoxia.** Frozen-thawed BDF2 morulae recovered at 2.5 DPC were cultured at 37°C under 5% $O_2$ and 5% $CO_2$, and treatment with 200 nM INK128 was initiated 24 h later at 3.5 DPC. Representative brightfield images are shown at 3.5 DPC (day 0 of treatment), 5.5 DPC (day 2), 10.5 DPC (day 7), and 19.5 DPC (day 16); the number of morphologically intact embryos of the 10 cultured is given above each panel. Embryos declined from 10 of 10 at treatment onset to 6 of 10 by day 2 and 3 of 10 by day 7, after which the surviving embryos remained intact and arrested through day 16. Inset, magnified view of two paused blastocysts at 19.5 DPC retaining defined blastocoel and inner cell mass structure.

Fig. S4

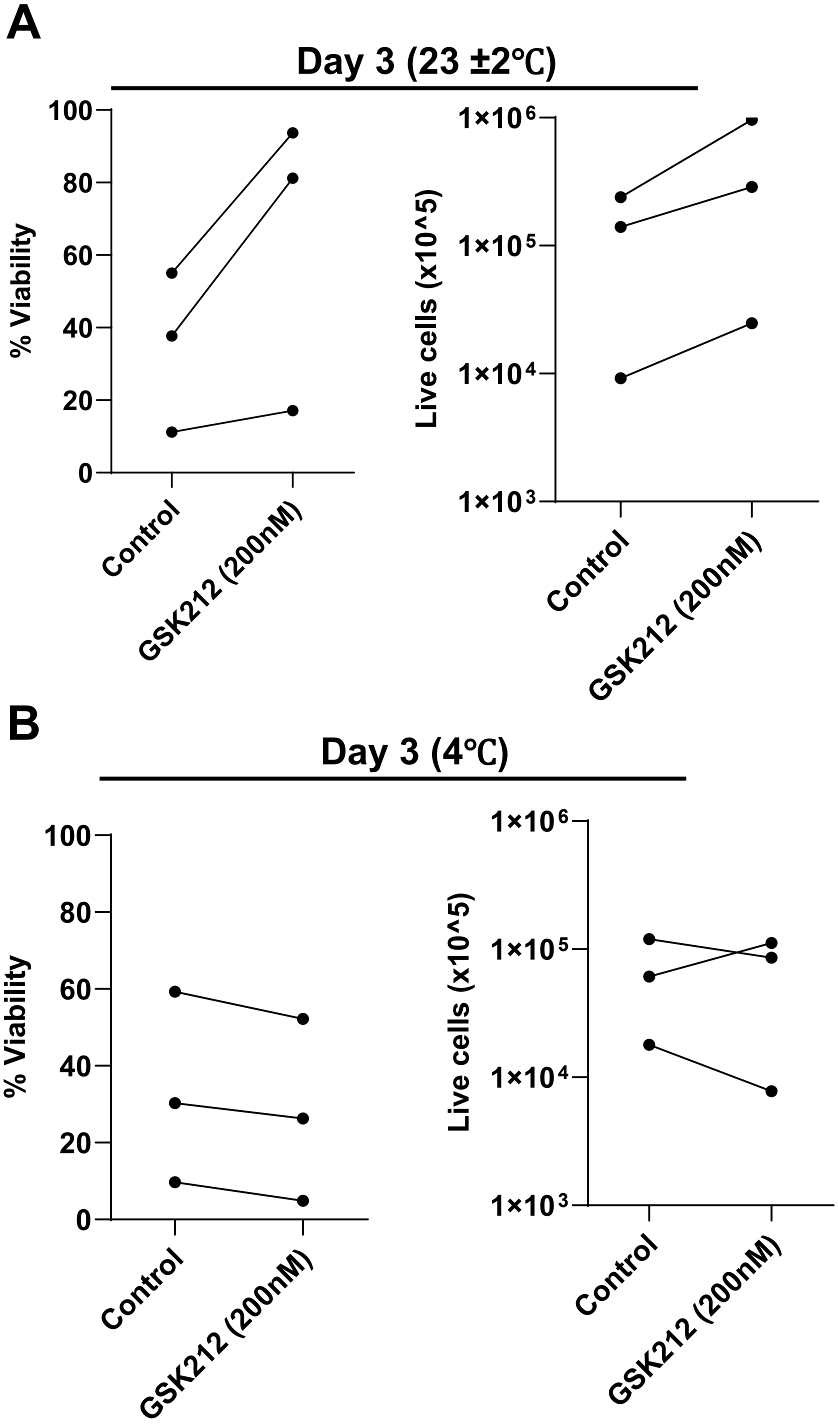


**Fig. S4. GSK2126458 preserves mESC viability during sub-normothermic but not deep-cold storage.** SGE2 mESCs (1 × $10^5$ per cryovial) were stored for 3 days in GMEM + 10% FBS + 2iL medium, without (Control) or with 200 nM GSK212, then assessed for recovery (n = 3 independent experiments; lines connect paired replicates). (**A**) At room temperature (23 ± 2°C), GSK212 yielded higher viability (left) and greater live-cell recovery (right) than untreated controls in all three replicates. (**B**) At 4°C, GSK212 conferred no such benefit, with viability and live-cell recovery comparable to or below control, consistent with reduced metabolic activity at deep cold limiting the pausing effect. Raw counts are provided in table S6.

Fig. S5

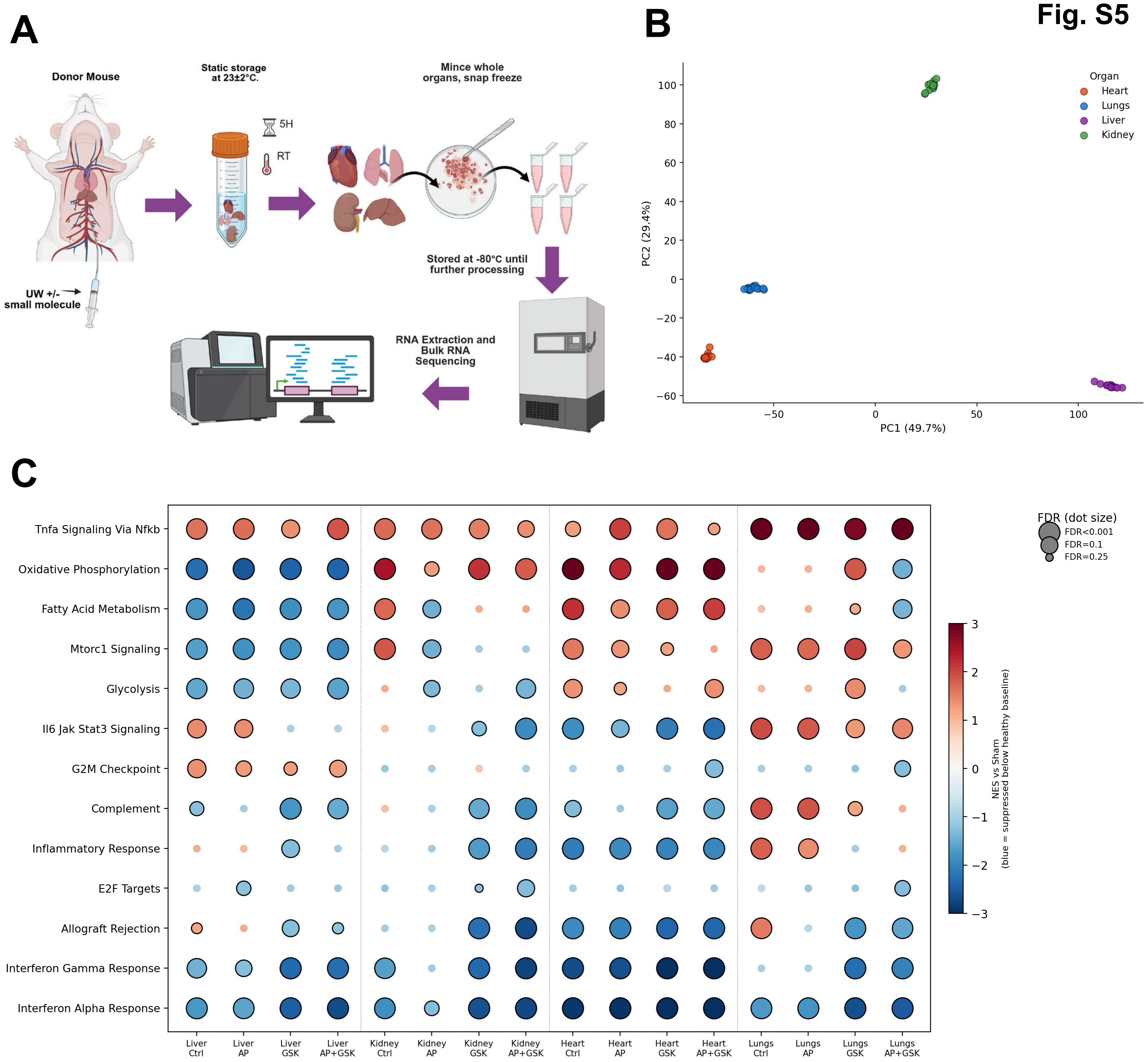


**Fig. S5. Whole-organ perfusion and bulk transcriptomic profiling of TorPause-treated mouse organs.** (**A**) Experimental workflow. Donor mice were perfused with University of Wisconsin (UW) solution alone (Control, 0.1% DMSO vehicle) or supplemented with small-molecule treatment (AP39 (3 µM), GSK212 (2 µM), or AP39 (3 µM) + GSK212 (2 µM)); organs were subjected to 5 hours of static storage at room temperature (23 ± 2°C), and whole heart, lung, liver, and kidney were minced, snap-frozen, and stored at −80°C prior to RNA extraction and stranded bulk RNA-sequencing. A separate Sham group was perfused and organs snap-frozen immediately, without storage, to provide a fresh, non-stored baseline. **(B)** Principal-component analysis of all samples (top 2,000 variable genes; n = 60). Samples separate by organ of origin, confirming data quality and sample identity. **(C)** Hallmark gene-set enrichment (GSEA prerank on DESeq2 Wald statistics; MSigDB Mouse Hallmark) for each condition relative to the Sham baseline, within organ. Color, normalized enrichment score (NES; blue, suppressed below the Sham baseline; red, elevated); dot size, false-discovery rate. Relative to Sham, GSK212 and the combination coordinately deepen suppression of interferon and inflammatory programs across organs, and suppress the allograft-rejection signature in kidney and lung, with metabolic programs (oxidative phosphorylation, mTORC1, glycolysis) suppressed predominantly in the liver, together indicating an induced torpor-like transcriptional state. n = 3 biological replicates per organ × condition. Full statistics in table S7.

Fig. S6

**A**

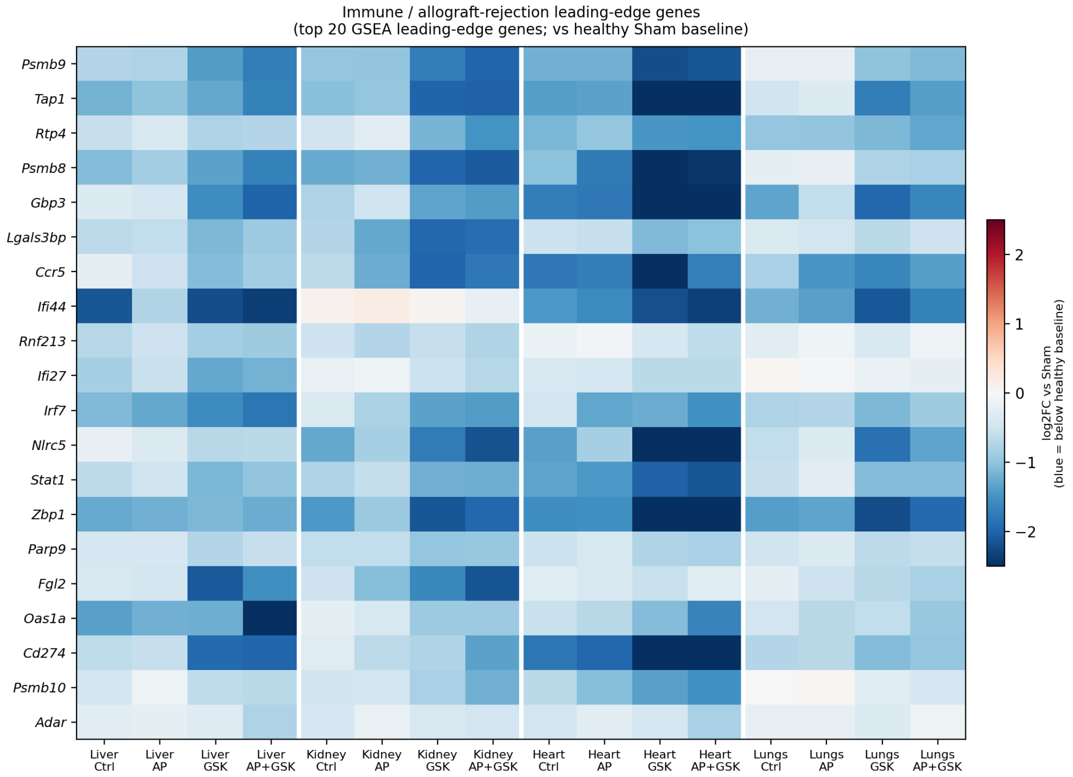

Immune / allograft-rejection leading-edge genes
(top 20 GSEA leading-edge genes; vs healthy Sham baseline)
Psmb9
Tap1
Rtp4
Psmb8
Gbp3
Lgals3bp
Ccr5
Ifi44
Rnf213
Ifi27
Irf7
Nlrc5
Stat1
Zbp1
Parp9
Fgl2
Oas1a
Cd274
Psmb10
Adar
Liver Ctrl
Liver AP
Liver GSK
Liver AP+GSK
Kidney Ctrl
Kidney AP
Kidney GSK
Kidney AP+GSK
Heart Ctrl
Heart AP
Heart GSK
Heart AP+GSK
Lungs Ctrl
Lungs AP
Lungs GSK
Lungs AP+GSK
log2FC vs Sham
(blue = below healthy baseline)
2
1
0
−1
−2


**B**

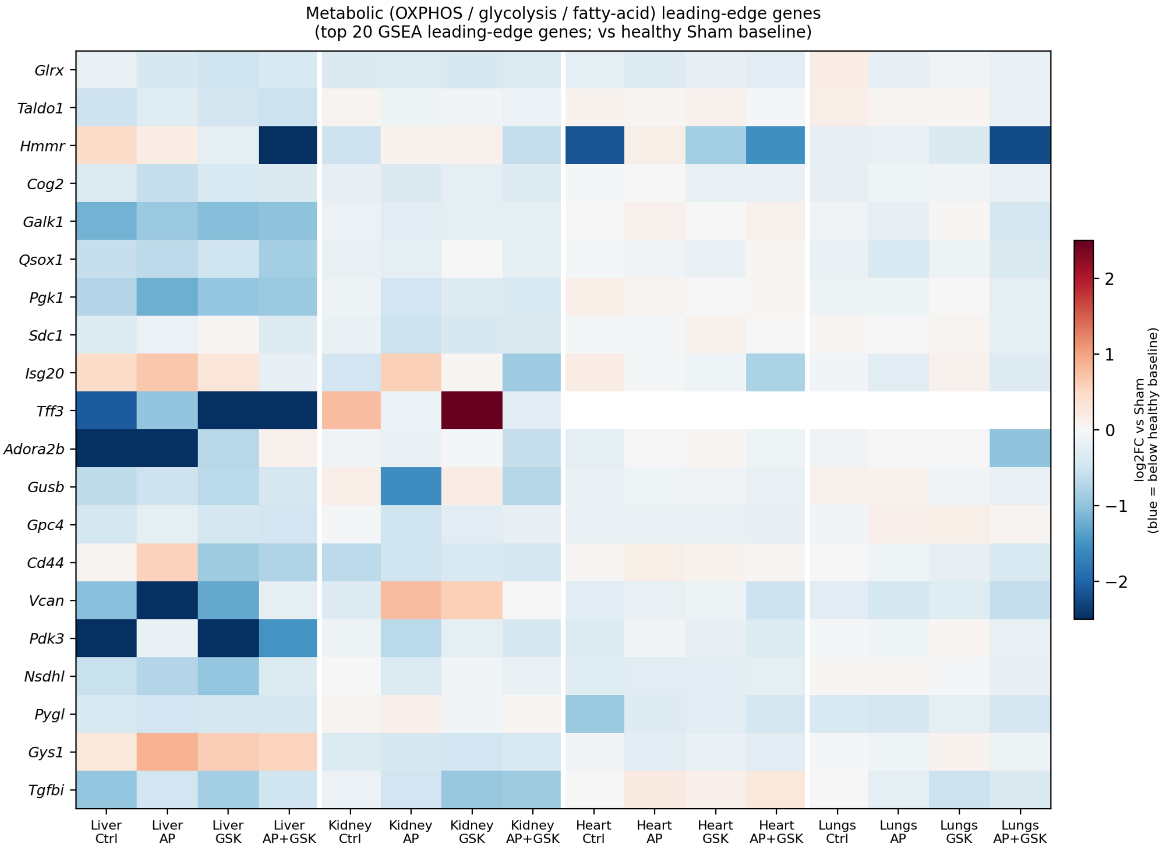

Metabolic (OXPHOS / glycolysis / fatty-acid) leading-edge genes
(top 20 GSEA leading-edge genes; vs healthy Sham baseline)
Glrx
Taldo1
Hmmr
Cog2
Galk1
Qsox1
Pgk1
Sdc1
Isg20
Tff3
Adora2b
Gusb
Gpc4
Cd44
Vcan
Pdk3
Nsdhl
Pygl
Gys1
Tgfbi
Liver Ctrl
Liver AP
Liver GSK
Liver AP+GSK
Kidney Ctrl
Kidney AP
Kidney GSK
Kidney AP+GSK
Heart Ctrl
Heart AP
Heart GSK
Heart AP+GSK
Lungs Ctrl
Lungs AP
Lungs GSK
Lungs AP+GSK
log2FC vs Sham
(blue = below healthy baseline)
2
1
0
−1
−2

**Fig. S6. Leading-edge genes of suppressed immune and metabolic programs across TorPause-treated organs.** Heat maps show per-gene log2 fold-change relative to the Sham baseline (blue, below baseline; red, above) for the top 20 leading-edge genes of the indicated Hallmark programs, defined by GSEA (genes contributing most to each enrichment, not selected by fold-change), for each organ and condition (Control, AP39, GSK212, AP39 + GSK212). (**A**) Immune and allograft-rejection leading-edge genes (pooled from the interferon-γ, interferon-α, and allograft-rejection Hallmark sets). GSK212 and the combination coordinately suppress antigen-processing (e.g., *Psmb8*, *Psmb9*, *Tap1*), interferon-regulatory (e.g., *Nlrc5*, *Irf7*, *Gbp3*), and rejection-associated genes below baseline across all four organs. (**B**) Metabolic leading-edge genes (pooled from the oxidative-phosphorylation, glycolysis, and fatty-acid-metabolism Hallmark sets). Suppression is predominant in the liver; direction is mixed in the other organs. n = 3 biological replicates per organ × condition; full gene lists in table S7.

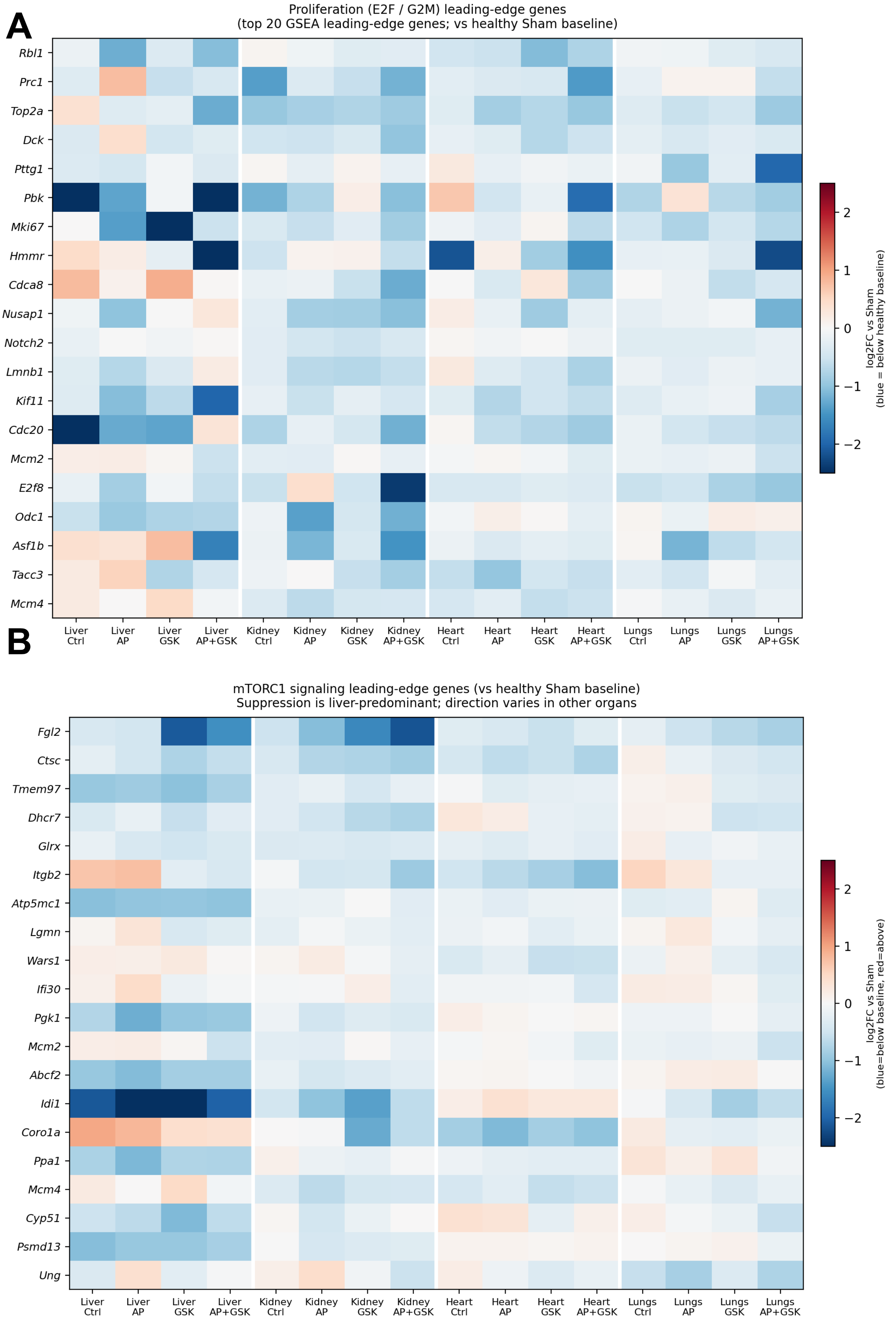
A
Proliferation (E2F / G2M) leading-edge genes
(top 20 GSEA leading-edge genes; vs healthy Sham baseline)
Rbl1
Prc1
Top2a
Dck
Pttg1
Pbk
Mki67
Hmmr
Cdca8
Nusap1
Notch2
Lmnb1
Kif11
Cdc20
Mcm2
E2f8
Odc1
Asf1b
Tacc3
Mcm4
Liver Ctrl
Liver AP
Liver GSK
Liver AP+GSK
Kidney Ctrl
Kidney AP
Kidney GSK
Kidney AP+GSK
Heart Ctrl
Heart AP
Heart GSK
Heart AP+GSK
Lungs Ctrl
Lungs AP
Lungs GSK
Lungs AP+GSK
2
1
0
−1
−2
log2FC vs Sham
(blue = below healthy baseline)
B
mTORC1 signaling leading-edge genes (vs healthy Sham baseline)
Suppression is liver-predominant; direction varies in other organs
Fgl2
Ctsc
Tmem97
Dhcr7
Glrx
Itgb2
Atp5mc1
Lgmn
Wars1
Ifi30
Pgk1
Mcm2
Abcf2
Idi1
Coro1a
Ppa1
Mcm4
Cyp51
Psmd13
Ung
Liver Ctrl
Liver AP
Liver GSK
Liver AP+GSK
Kidney Ctrl
Kidney AP
Kidney GSK
Kidney AP+GSK
Heart Ctrl
Heart AP
Heart GSK
Heart AP+GSK
Lungs Ctrl
Lungs AP
Lungs GSK
Lungs AP+GSK
2
1
0
−1
−2
log2FC vs Sham
(blue=below baseline, red=above)

**Fig. S7**

**Fig. S7. Leading-edge genes of proliferative and mTORC1-signaling programs across TorPause-treated organs.** Heat maps show per-gene log2 fold-change relative to the Sham baseline (blue, below baseline; red, above) for the top 20 GSEA leading-edge genes of the indicated Hallmark programs, for each organ and condition. (**A**) Proliferation leading-edge genes (pooled from the E2F-targets and G2M-checkpoint Hallmark sets), including *Mki67*, *Top2a*, *Cdc20*, and *Kif11*. Suppression is modest and variable across organs, most evident in the liver. (**B**) mTORC1-signaling leading-edge genes. Suppression is liver-predominant, with mixed direction in the other organs. n = 3 biological replicates per organ × condition; full gene lists in table S7.

**Fig S8**

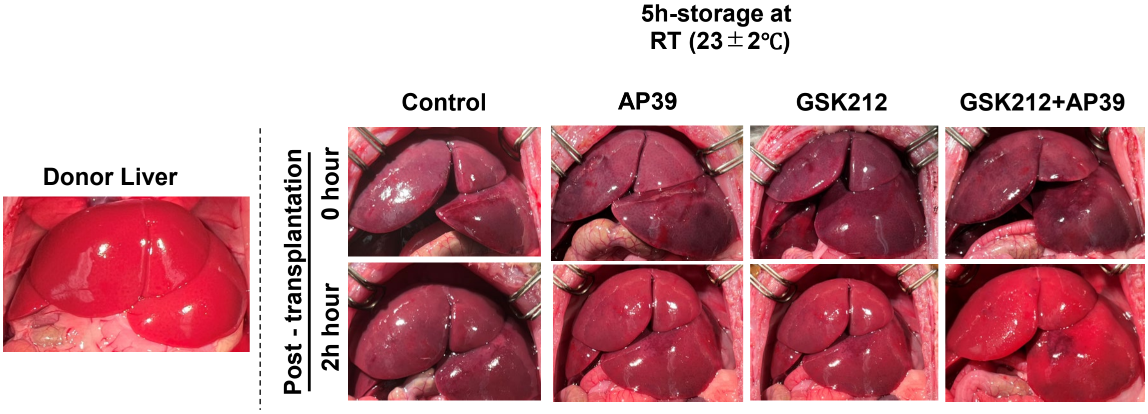


**Fig. S8. Macroscopic evaluation of liver grafts post-transplantation.** Representative gross morphological images of rat livers at reperfusion (0 hour) and 2 hours post-transplantation following 5 hours of ambient room-temperature (23 ± 2°C) storage in preservation solution alone (Control) or supplemented with AP39, GSK212, or GSK212 + AP39 (n = 3 per group). A representative freshly procured donor liver is shown at left for reference. The combinatorial treatment (GSK212 + AP39) restored and maintained healthy graft coloration compared with the severe congestion seen in untreated controls.

**Fig. S9**

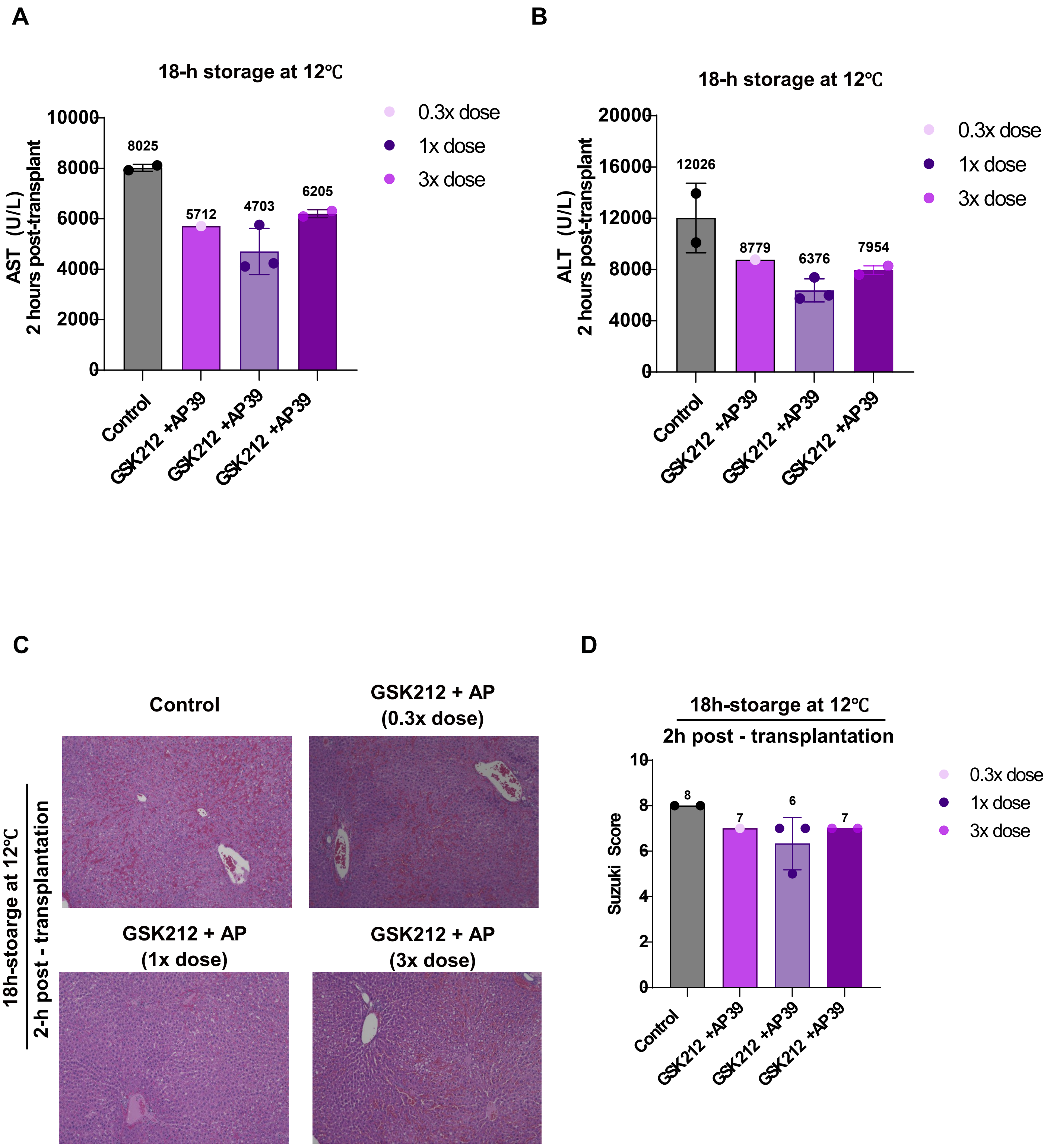

A
18-h storage at 12℃
AST (U/L)
2 hours post-transplant
10000
8000
6000
4000
2000
0
8025
5712
4703
6205
Control
GSK212 +AP39
GSK212 +AP39
GSK212 +AP39
0.3x dose
1x dose
3x dose
B
18-h storage at 12℃
ALT (U/L)
2 hours post-transplant
20000
16000
12000
8000
4000
0
12026
8779
6376
7954
Control
GSK212 +AP39
GSK212 +AP39
GSK212 +AP39
0.3x dose
1x dose
3x dose
C
Control
GSK212 + AP
(0.3x dose)
GSK212 + AP
(1x dose)
GSK212 + AP
(3x dose)
18h-stoarge at 12℃
2-h post - transplantation
D
18h-stoarge at 12℃
2h post - transplantation
Suzuki Score
10
8
6
4
2
0
8
7
6
7
Control
GSK212 +AP39
GSK212 +AP39
GSK212 +AP39
0.3x dose
1x dose
3x dose

**Fig. S9 Dose-ranging of combined GSK212 + AP39 treatment following 18-hour storage at 12°C.** Rat livers were stored for 18 hours at 12°C in preservation solution alone (Control) or supplemented with GSK212 + AP39 at 0.3×, 1×, or 3× the reference concentration, then orthotopically transplanted. At 1×, the perfusate contained 2 μM GSK212 and 3 μM AP39 with a 600 nM AP39 bolus; all components were scaled together by the indicated multiplier. (**A and B)** Serum AST (A) and ALT (B) 2 hours post-transplantation. (**C**) Representative hematoxylin and eosin–stained graft sections collected 2 hours post-transplantation. (**D**) Suzuki injury score for the sections in **(C)**. Bars show the group mean and each symbol represents one animal; the value above each bar is the group mean, and error bars show SD where n ≥ 2 (Control, n = 2; 0.3×, n = 1; 1×, n = 3; 3×, n = 2).

**Fig. S10**

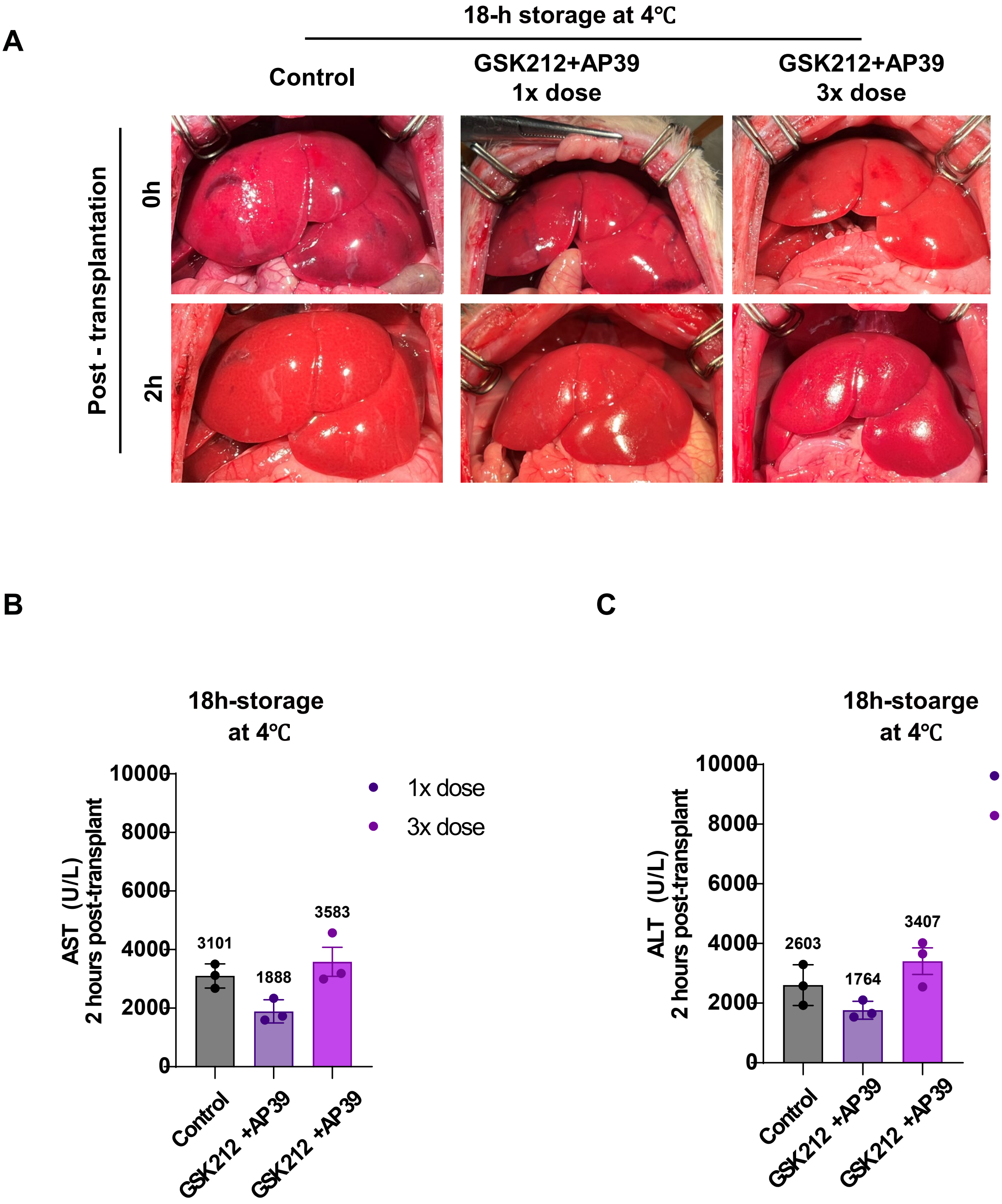


**Fig. S10. Dose-ranging of combined GSK212 + AP39 treatment following 18-hour storage at 4°C.** Rat livers were stored for 18 hours at 4°C in preservation solution alone (Control) or supplemented with GSK212 + AP39 at 1× or 3× the reference concentration, then orthotopically transplanted. At 1×, the perfusate contained 2 μM GSK212 and 3 μM AP39 with a 600 nM AP39 bolus; all components were scaled together by the indicated multiplier. (**A**) Representative gross appearance of the graft immediately after reperfusion (0 hour) and at 2 hours post-transplantation. (**B** and **C**) Serum AST (B) and ALT (C) 2 hours post-transplantation (n = 3 animals per group). Bars show group mean ± SD; each symbol represents one animal, and the value above each bar is the group mean. The 1× dose produced significantly lower transaminase release than the 3× dose (AST, $P$ = 0.030; ALT, $P$ = 0.040; one-way ANOVA with post hoc pairwise comparison). Full statistical details are reported in table S9.
